# Extracellular Vesicles Enable CircRNA Delivery via in situ Biogenesis and Sorting

**DOI:** 10.64898/2026.04.24.720573

**Authors:** Minchao Li, Yuxuan Pan, Mingting Cui, Junyao Deng, Fei Wang, Lanya Li, Rui Zhang, Caijun Sun, Zhenhua Li

## Abstract

Extracellular vesicles (EVs) are promising vehicles for nucleic acid delivery, yet efficient delivery of circular RNA (circRNA) remains challenging due to inefficient loading and limited intracellular expression. Here, we establish an EV-based platform that enables efficient circRNA delivery via in situ biogenesis and sorting. By optimizing intracellular circularization and translation through vector design, we markedly enhance circRNA expression. By combining this with Snu13-mediated EV sorting and enhanced vesicle biogenesis, we achieve efficient packaging of circRNA without compromising vesicle integrity. This integrated strategy enables robust and sustained protein expression following EV-based circRNA delivery. By leveraging this platform, we demonstrate a dendritic cell-targeting circRNA vaccine that elicits strong antigen-specific CD8^+^ T cell responses and antitumor efficacy. We further show that systemic delivery of BNP-encoding circRNA attenuates doxorubicin-induced myocardial fibrosis. Together, this work establishes a generalizable platform for circRNA therapeutics by overcoming key barriers in circRNA expression and EV-mediated delivery.

**Teaser:** Engineered EVs enable efficient circRNA delivery for sustained protein expression and therapy.

## Introduction

Extracellular vesicles (EVs) have emerged as promising vehicles for nucleic acid delivery owing to their intrinsic biocompatibility, low immunogenicity, and efficient cellular uptake(*1–3*). Importantly, compared to lipid nanoparticles (LNPs), EVs exhibit more efficient cellular uptake, driven by their nanoscale architecture and the presence of endogenous membrane proteins (such as tetraspanins and integrins) that facilitate cell recognition and internalization(*4, 5*). Nevertheless, the clinical potential of EV-mediated delivery of coding RNA, including messenger RNA (mRNA), microRNA (miRNA), and circular RNA (circRNA), remains constrained by inefficient cargo loading, limiting both functional studies and therapeutic applications.

Therapeutic messenger RNAs (mRNAs) have found broad applications in vaccine development and the treatment of genetic disorders, particularly owing to the remarkable success of mRNA-based COVID-19 vaccines. Nevertheless, conventional linear mRNAs carry terminal motifs that are recognized by exonucleases, rendering them prone to rapid degradation and consequently conferring a short half-life in physiological environments. Moreover, these mRNAs can engage various pattern recognition receptors, leading to undesired innate immune responses. While conventional mRNA therapies are hampered by the short cytoplasmic half-life of linear transcripts, circRNA offers a compelling alternative. Its covalently closed structure confers superior resistance to nucleases, enabling prolonged protein expression in vivo(*6, 7*). Translation of circRNA occurs in a cap-independent manner via internal ribosome entry sites (IRES), and large-scale production can be achieved through enzymatic cyclization or self-splicing intron systems(*8*). In addition, the low immunogenicity of circRNA endows them with a marked benefit for therapeutic applications.

Despite these advantages, efficient delivery of circRNA via EVs remains challenging. Typically, the loading of therapeutic RNAs into EVs can be achieved through two types of methods. The first type introduces RNAs after EV purification, exemplified by electroporation and sonication. Such post-isolation loading techniques, however, tend to compromise the structural integrity of EVs and may also damage the RNA molecules themselves(*9*). A different approach entails cargo encapsulation before EV purification, which takes advantage of the natural sorting processes occurring within the donor cells. However, this method often exhibits inadequate loading efficiency, and its mechanistic basis is still unclear. The endogenous loading approaches preserve vesicle structure and support the incorporation of larger RNA cargos, but are limited by inherently low intracellular circRNA expression, with transcript levels generated by endogenous processing elements remaining substantially lower than those of linear mRNAs(*10, 11*). Moreover, even when expression is enhanced, passive enrichment of circRNA into EVs remains inefficient. Together, these limitations represent a critical barrier to efficient circRNA delivery via EVs, necessitating strategies that simultaneously enhance circRNA expression and EV loading.

To address these limitations, we developed a multi-pronged strategy for efficient EV-mediated circRNA delivery. First, we designed a circRNA expression vector based on a tRNA cleavage-mediated cyclization mechanism and systematically optimized its sequence to enhance both transcriptional and translation efficiency in mammalian cells. Second, we engineered the EV membrane by displaying an RNA-binding protein capable of recognizing and enriching circRNA within vesicles. Third, we augmented EV biogenesis and secretion through the co-expression of key regulatory genes. By integrating these approaches, we established a robust platform for high-yield circRNA encapsulation and functional delivery, as validated through in vivo studies (**Fig. 1**). Using this platform, we further demonstrated the versatility of EV-mediated circRNA delivery across circRNA cargos with distinct lengths and biological functions, including a dendritic cell-targeting circRNA vaccine that elicited strong antigen-specific CD8⁺ T cell responses and antitumor efficacy, as well as a BNP-encoding circRNA therapeutic that attenuated doxorubicin-induced myocardial fibrosis. Collectively, this work provides a scalable and efficient engineering framework for EV-based circRNA therapeutics, offering new avenues for RNA delivery.

**Fig. 1.**
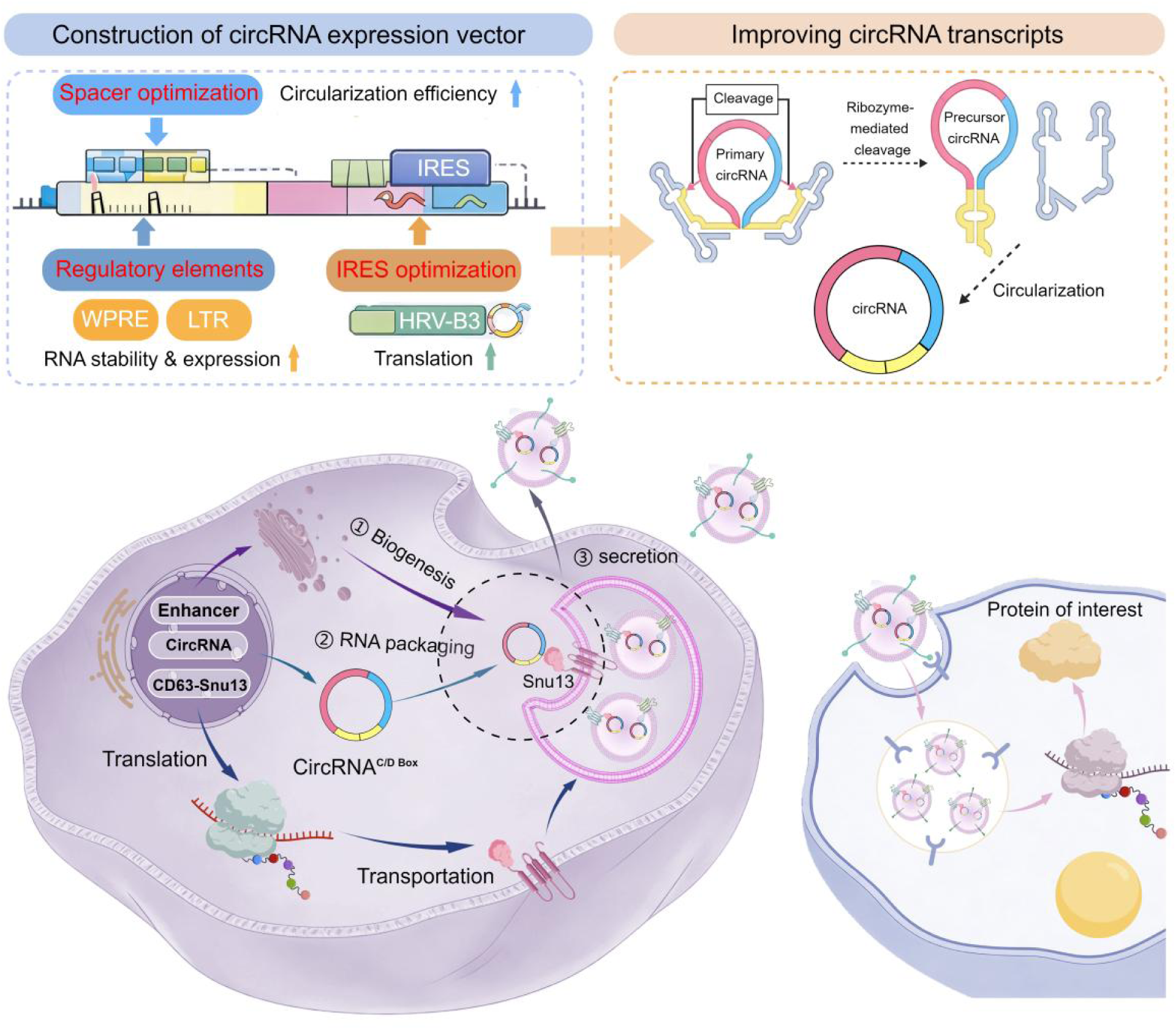
Schematic of circRNA expression vector construction and EV loading. The diagram illustrates the construction of an optimized circRNA expression vector containing specific spacers, regulatory elements (WPRE, LTR), and IRES sequences to ensure stability and translation. Following transcription and circularization, the circRNA (containing a C/D box) is specifically recruited to EVs via interaction with the CD63-Snu13 fusion protein. This mechanism promotes efficient in situ packaging and secretion of circRNA. In parallel, enhancement of EV biogenesis and secretion through co-expression of key regulatory factors (STEAP3, SDC4, and NadB) increased EV output and overall circRNA payload. Finally, the engineered EVs deliver the circRNA to recipient cells for the functional production of the target protein.

## Results

### Optimized circRNA Expression Vectors Enable Efficient Transcription and Translation

CircRNA can be produced in mammalian cells from a plasmid vector using a ribozyme-based self-cleavage system. This approach involves transcribing a linear RNA with ribozymes encoded at its 5’ and 3’ ends. Following self-catalyzed cleavage by the ribozymes, the resulting ends are ligated by the endogenous RNA ligase RtcB to form circular RNA(*11*). We designed a linear RNA transcript driven by the CMV promoter, which encoded the 5’ UTR, IRES, Kozak sequence, coding sequence (CDS), C/D Box motif, and 3’ UTR between twister ribozyme sequences (**Fig. 2A**). The transcribed ribozymes underwent rapid self-cleavage, thereby exposing complementary ligation stems that facilitated end joining and subsequent in situ circularization. Correct circularization was confirmed by Sanger sequencing across the junction site (**Fig. 2B**).

**Fig. 2.**
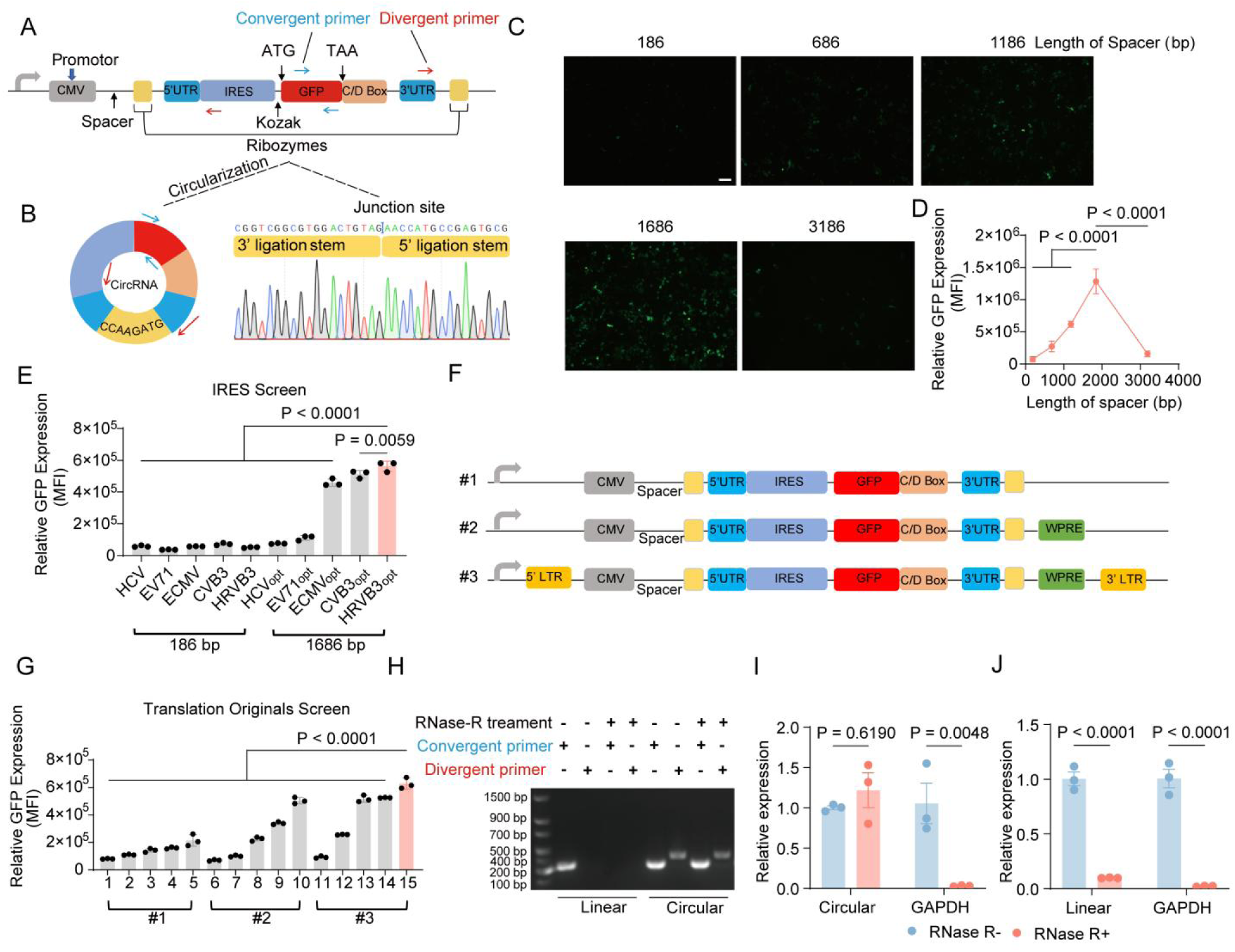
Development and Characterization of an Optimized Intracellular Expression Vector for CircRNA. (A) Schematic of plasmid DNA for the production of circRNA. The circRNA are produced through transfecting HEK-293T cells to generate linear RNA precursors. Each precursor is designed with an IRES-coupled mRNA of interest flanked by twister ribozymes. Following transcription, the ribozymes undergo rapid self-cleavage, which exposes complementary ligation stems and allows them to hybridize. Subsequently, these pre-linear RNAs serve as substrates for the ubiquitous endogenous RNA ligase RtcB, which catalyzes their circularization in situ. (B) The characteristic back-splicing of the circRNA validated by Sanger sequencing of the PCR-amplified junction. (C) Length-dependent enhancement of GFP expression by engineered spacer sequences. The HEK-293T cells were treated with different circRNA expression vector insert various length of spacer. Fluorescence signal using inverted fluorescence microscope and the GFP expression efficiency was quantified by (D) flow cytometry. Scale bar represents 200 μm. (E) The performance of diverse IRES sequences within the circRNA expression vector was assessed in HEK-293T cells. Following transfection, GFP intensity was measured quantitatively via flow cytometry. (F) Schematic of the engineered circRNA expression plasmid. The vector architecture includes the WPRE and LTR element to enhance translational efficiency. (G) The performance of WPRE and the LTR element within the circRNA expression vector was assessed in HEK-293T cells. The GFP intensity was measured quantitatively via flow cytometry. Three vector sets (#1: plasmids 1-5; #2: plasmids 6-10; #3: plasmids 11-15) were constructed, each containing the same series of IRES elements in the order of HCV, EV71, ECMV, CVB3, and HRV-B3. (H) The successful intracellular expression of circRNA was confirmed by RT-PCR using divergent primers, which produced a specific amplification product for circRNA^GFP^ from cDNA but not from linear mRNA controls. The circular structure of circRNA^GFP^ was verified by its resistance to RNase R exonuclease digestion. (I-J) The differential susceptibility of circRNA^GFP^ and mRNA^GFP^ to RNase R exonuclease was evaluated by RT-qPCR. Data are presented as mean ± SD. Statistical analysis in D was using two-tailed unpaired Student’s t-tes. Statistical analysis in E and G were using one-way ANOVA with a Tukey multiple comparisons test. Statistical analysis in I and J were using two-way ANOVA with a Tukey multiple comparisons test.

To enhance the transcription and translation efficiency of circRNA expression vectors, we performed sequence optimization. Spacer insertion between the CMV promoter and twister ribozymes modulated circRNA expression in a length-dependent manner, with a 1,686 bp spacer achieving maximal GFP expression (**Fig. 2C-D and fig. S1A**). Quantitative analysis revealed that this optimized vector yielded an approximately 3000-fold increase in circRNA^GFP^ transcript levels compared to the unmodified construct (**fig. S1B**). However, a further increase in spacer length led to a decline in both transcription and expression, indicating that while spacer insertion is beneficial, excessive template length is detrimental to efficiency. To further enhance the translational efficiency of circRNA, we screened five natural and synthetic internal ribosome entry site (IRES) elements(*10, 12*) (**Table S1**). The IRES from human rhinovirus B3 (HRV-B3) demonstrated superior performance, facilitating the highest level of cap-independent translation. This efficiency was further augmented by the incorporation of spacer sequences (**Fig. 2E and fig. S1C**).

Since the tRNA-based cyclization system involves cleavage by twister ribozymes, resulting in the loss of the 5’ cap and 3’ poly(A) tail, any linear RNA is rapidly degraded upon cytoplasmic export, thereby reducing its abundance(*13*). To counter this, we incorporated the WPRE, which promotes nuclear export of RNA and enhances transcript stability(*14*), thereby increasing circRNA yield. Furthermore, the addition of a long terminal repeat (LTR) element, which exhibits enhancer activity, further augmented the translational efficiency of the circRNA expression construct (**Fig. 2F and Table S2**). The inclusion of WPRE increased GFP expression by approximately 2.5-fold. A further enhancement, achieving an approximately 3-fold increase in GFP expression, was observed when both WPRE and the LTR element were incorporated (**Fig. 2G and fig. S1D**). Consequently, all subsequent experiments utilized the optimized vector comprising HRV-B3 IRES, WPRE, and LTR components.

To molecularly characterize the intracellular transcripts, we performed reverse transcription-PCR (RT-PCR) using primers flanking the predicted back-splice junction. Agarose gel analysis confirmed the specific amplification of circRNA^GFP^ from cDNA, which was absent in control samples of linear mRNA (**Fig. 2H**). Corroborating these findings, an RNase R digestion assay was conducted. This exonuclease selectively degrades linear RNAs, and results showed that circRNA^GFP^ remained largely intact while mRNA^GFP^ was rapidly degraded, confirming its circular topology and enhanced inherent stability (**Fig. 2I-J**).

### EV–based delivery of circular RNA via in situ biogenesis and sorting

To establish a robust EV platform for efficient circRNA delivery, we implemented a dual-strategy approach to enhance both circRNA encapsulation and EV production. Specifically, we engineered the EV membrane to display RNA-binding proteins that selectively enrich circRNA within vesicles. In parallel, we enhanced cellular pathways governing EV biogenesis and secretion to increase EV yield, thereby elevating the total circRNA payload **(Fig. 3A)**. Based on prior evidence that the RNA-binding protein Snu13 specifically recognizes the K-turn structural motif within C/D Box RNA sequences to form ribonucleoprotein complexes(*15*), we fused Snu13 to the C-terminus of the tetraspanin CD63 (a canonical EV membrane marker). Overexpression of the CD63-Snu13 fusion protein enabled the display of Snu13 on the luminal side of EV membranes. Concurrently, we inserted a C/D Box motif downstream of the stop codon in a circRNA^GFP^ expression construct to prevent any interference with proper protein translation. Co-transfection of HEK-293T cells with plasmids encoding circRNA^GFP-C/D Box^ and CD63-Snu13 facilitated the assembly of circRNA-protein complexes inside EVs. RNA immunoprecipitation (RIP) assays confirmed the specific interaction between Snu13 and the C/D Box-tagged circRNA, as RT-qPCR analysis demonstrated significant enrichment of circRNA in CD63-Snu13 pulldowns **(Fig. 3B and fig. S2A)**. To boost EV biosynthesis, we designed a synthetic polyprotein construct integrating STEAP3 (a regulator of endosomal vesicle formation)(*16*), syndecan-4 (SDC4; promotes intraluminal vesicle budding)(*17*), and a fragment of L-aspartate oxidase (NadB; modulates tricarboxylic acid cycle activity to potentially enhance metabolic fitness)(*18*). We also constructed a CD63-nanoluciferase (CD63-Nluc) reporter to quantify EV secretion. HEK-293T was co-transfected with the biogenesis enhancing construct and CD63-Nluc plasmid, and following removal of free Nluc in culture supernatants via centrifugation. Subsequently, the luminescence intensity of the cell culture supernatant was detected. We found that the combined expression of these genes significantly increased EV production **(Fig. 3C)**. Nanoparticle tracking analysis (NTA) further confirmed a significant increase in EV particle concentration without altering the typical size distribution profile **(fig. S2B)**.

**Fig. 3.**
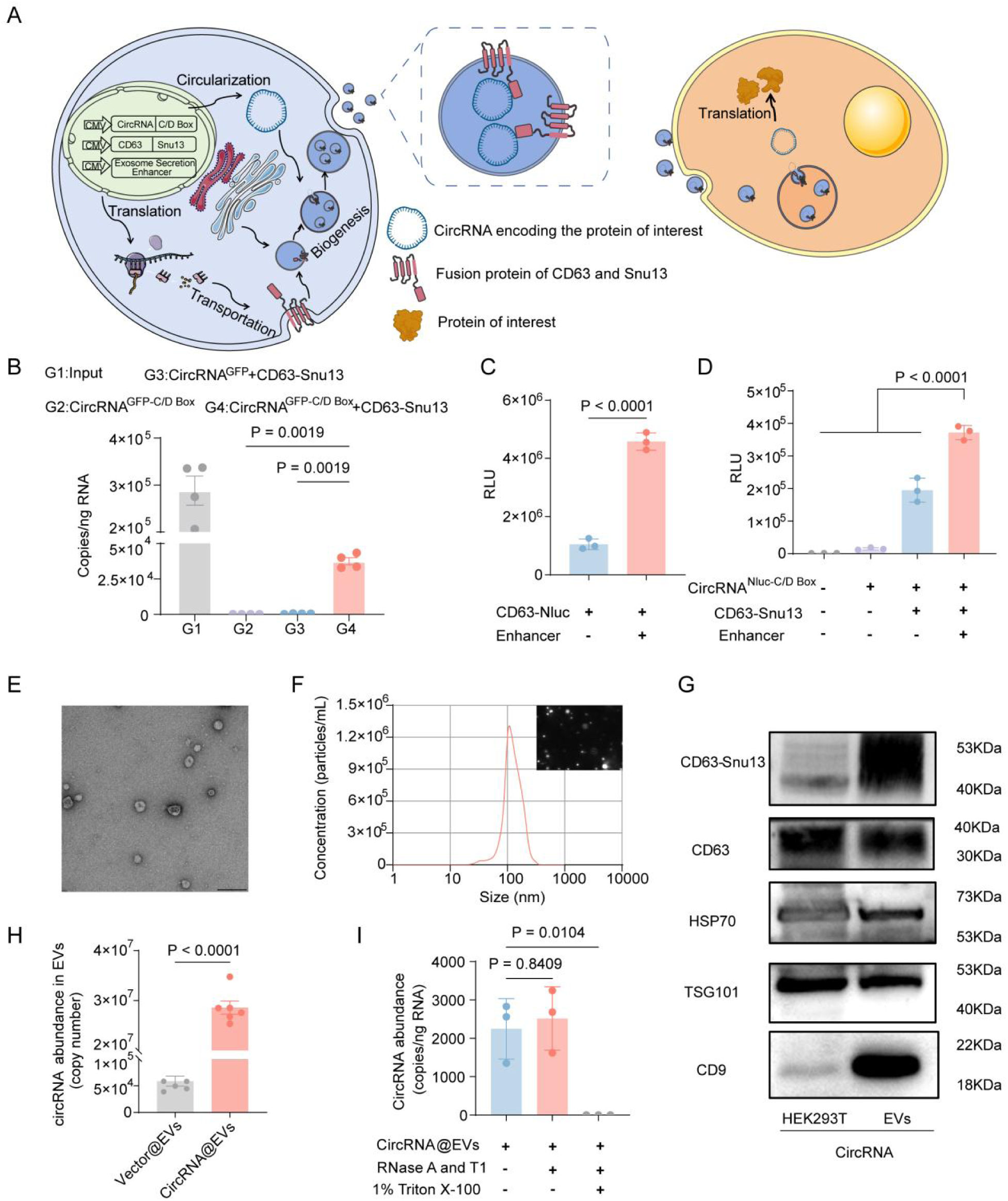
Construction and characterization of circRNA-Loaded Extracellular Vesicles. (A) Schematic illustration of the engineered EV platform for circRNA delivery. Producer cells were co-transfected with an exosome production enhancer to boost EV biogenesis, along with constructs encoding an RNA-packaging device (CD63–Snu13) and a C/D box–tagged circRNA (e.g., circRNA^Nluc^). The resulting engineered EVs, loaded with circRNA, were isolated and applied to recipient cells (HEK293T or BMDCs). Upon internalization, the circRNA was translated into functional protein (e.g., Nluc) in the target cells. (B) RIP analysis demonstrating specific interaction between CD63-Snu13 and C/D box-containing circRNA^GFP^ using anti-HA antibody. (C) Enhanced EVs secretion was evaluated by measuring Nluc activity in supernatants following sequential centrifugation to remove cells and debris. RLU represents Relative Light Unit. (D) Functional delivery of circRNA via engineered EVs. HEK-293T cells were transfected with plasmids encoding circRNA^Nluc-C/D^ ^Box^, EV production enhancer, and CD63-Snu13. Supernatants containing EVs were transferred to recipient cells, and Nluc activity was measured after 24 h. (E) Transmission electron micrograph of purified circRNA@EVs. Scale bar: 200 nm. (F) Size distribution profile of circRNA@EVs determined by nanoparticle tracking analysis (NTA). (G) Western blot analysis of EV markers (CD9, TSG101, HSP70, CD63) and CD63-Snu13 (HA-tagged) in cell lysates and purified EVs. (H) Absolute qPCR quantification of circRNA copies in engineered circRNA@EVs versus control EVs. (I) circRNA levels in circRNA@EVs treated with RNase A/T1 with or without Triton X-100 disruption. Data are presented as mean ± SD. Statistical analysis were using one-way ANOVA with a Tukey multiple comparisons test.

To evaluate the functionality of our engineered EV system, HEK-293T cells were co-transfected with three plasmids: a circRNA reporter encoding nanoluciferase (Nluc) containing the C/D Box motif, an RNA-enrichment construct (CD63-Snu13), and an EV biogenesis enhancer (STEAP3-SDC4-NadB). The culture supernatant containing secreted EVs was collected and applied directly to HEK-293T recipient cells. A pronounced luminescence signal was detected in recipient cells only when all three plasmids were co-expressed in the producer cells, indicating efficient packaging and functional delivery of circRNA^Nluc^ **(Fig. 3D)**. No signal was observed using supernatant from cells transfected with the circRNA vector alone. We also evaluated the circRNA enrichment capacity of the 1 × Snu13 versus the 3 × Snu13 protein and found the 1 × Snu13 version to be markedly more potent **(fig. S2C).** We next isolated and concentrated EVs from the supernatant and used bone marrow-derived dendritic cells (BMDCs) as recipients. Similarly, significant luminescence was measured in BMDCs **(fig. S2D)**, confirming that EV-delivered circRNA^Nluc^ can be functionally transferred into immune cells. For biophysical characterization, purified EVs were analyzed using transmission electron microscopy (TEM), nanoparticle tracking analysis (NTA), and western blot. TEM revealed that circRNA@EVs exhibited a uniform spherical morphology **(Fig. 3E)**. NTA indicated a peak particle diameter of 110.5 nm **(Fig. 3F)**. Western blot further confirmed the presence of canonical EV markers (CD9, TSG101, HSP70, CD63) as well as the engineered fusion protein CD63-Snu13 **(Fig. 3G)**, validating the identity and integrity of the vesicles. To quantify cargo loading, absolute qPCR was performed to determine circRNA copy number in EVs. We found that transfecting cells with the circRNA vector alone yielded only 5 × 10^4^ circRNA copies per 1 × 10^9^ EVs, whereas co-transfection with all three plasmids dramatically increased the yield to 3 × 10^7^ copies **(Fig. 3H)**, representing a significant enhancement in packaging efficiency. Finally, we tested the protective capacity of EVs for circRNA. Treatment of intact circRNA@EVs with an RNase A/T1 mixture did not reduce circRNA levels, whereas concurrent disruption of the EV membrane with 1% Triton X-100 led to complete degradation **(Fig. 3I)**, confirming that EVs effectively shield circRNA from enzymatic degradation.

In summary, we have developed a highly efficient and scalable EV-based platform for encapsulating and delivering functional circRNA. These engineered EVs not only protect circRNA from ribonuclease activity but also enable its functional delivery into multiple cell types, including HEK-293T cells and BMDCs.

### EVs-Based circRNA Vaccine Elicits a Potent Immune Response in Vitro and In Vivo

Leveraging the efficient encapsulation and delivery capabilities of EVs, we developed a vaccine platform for the delivery of ovalbumin (OVA)-encoding circRNA. We assessed protein expression from EVs loaded with circRNA encoding OVA in HEK-293T cells by western blot **(fig. S3A)**. To enhance vaccine antigen uptake by antigen-presenting cells (APCs), we engineered a DC-targeting strategy inspired by the previously reported RVG-LAMP2B fusion protein(*19*). Specifically, the DCpep sequence was cloned into the N-terminus of the LAMP2B gene to confer DC-specific targeting. Additionally, the amino acid sequence GNSTM was fused to the N-terminus of the resulting DCpep-Lamp2b construct to protect the chimeric protein from proteolytic degradation. The plasmid encoding GNSTM-DCpep-Lamp2b **(fig. S3B)** was then co-transfected into HEK-293T cells together with the RNA packaging device (CD63-Snu13), a C/D box-tagged circRNA, and an exosome production enhancer. EVs (circRNA^OVA^-DCpep@EVs) were subsequently isolated from the culture medium via density gradient centrifugation. We first assessed the innate immune activation and antigen presentation kinetics induced by this platform in bone marrow-derived dendritic cells (BMDCs) **(Fig. 4A)**. The uptake efficiency of PKH67-labeled circRNA^OVA^-DCpep@EVs by BMDCs was determined at different time points. Following incubation, cells were harvested and analyzed via flow cytometry to quantify EV internalization. As illustrated in **fig. S3C-D**, BMDCs exhibited time-dependent uptake of circRNA^OVA^-DCpep@EVs, reaching peak efficiency by 8 hours. We next investigated the capacity of circRNA^OVA^-DCpep@EVs to promote BMDC maturation. Compared to control treatments, all EV-based formulations significantly enhanced BMDC maturation, as indicated by the upregulated surface expression of the co-stimulatory molecules CD40 and CD86, as well as MHC I (**Fig. 4B and fig. S3E**). Furthermore, RT-qPCR analysis revealed that circRNA^OVA^-DCpep@EVs stimulated BMDCs to express a panel of pro-inflammatory cytokines and mediators, including IFN-γ, IL-6, IL-12, and TNF-α (**fig. S3F**). This enhanced immunostimulatory profile was corroborated by a multiplex cytokine assay, which confirmed the robust secretion of Th1-polarizing cytokines, known to be critical for anti-tumor immunity (**Fig. 4C and fig. S4**). Importantly, DCpep modification on the EV membrane consistently augmented DC activation across all readouts, highlighting its role as an effective targeting and immunostimulatory moiety. Successful antigen processing and presentation were subsequently evaluated by monitoring the surface display of the MHC I-OVA complex. As shown in **Fig. 4D and fig. S5**, the expression level of MHC I-OVA peaked at 6 hours post-treatment. Notably, the circRNA^OVA^-DCpep@EVs formulation sustained antigen presentation significantly longer than the non-targeted control (circRNA^OVA^@EVs) over a 48-hour period.

**Fig. 4.**
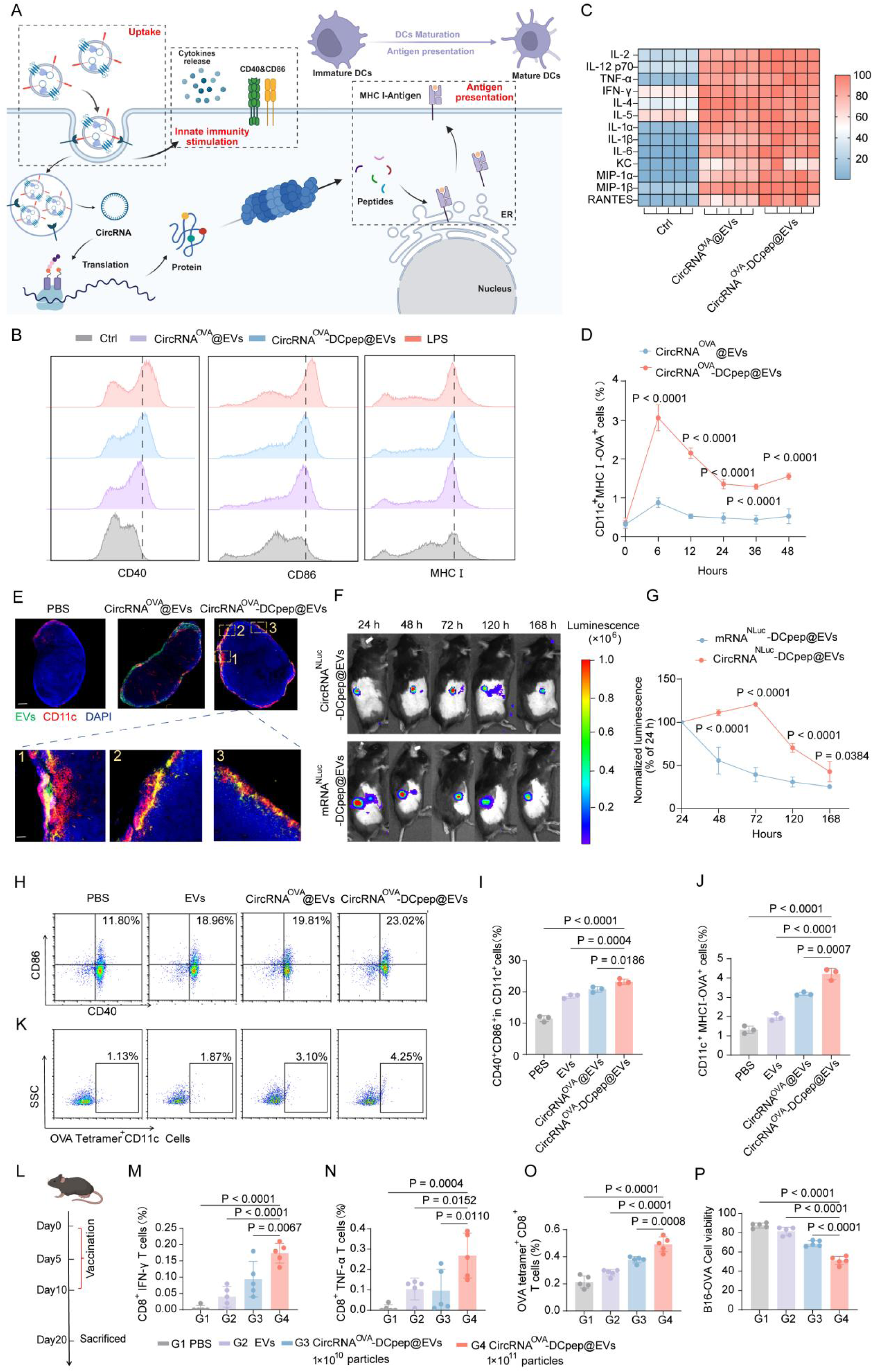
Immune response induced by the EVs-based circRNA vaccine. (A) Schematic illustration of the EV-based circRNA vaccine triggering innate immunity stimulation and antigen presentation. (B) Surface expression of maturation markers (CD40, CD86, MHC I) on CD11c^+^ BMDCs after exposure to different formulations. (C) Heatmap depicting the secretion profile of inflammatory cytokines by BMDCs incubated with different formulations for 24 h, analyzed using a multiplex immunoassay. (D) Kinetics of MHC I-OVA complex expression on BMDCs treated with circRNA^OVA^@EVs or circRNA^OVA^-DCpep@EVs, quantified by flow cytometry (n=3). (E) C57BL/6 mice were injected subcutaneously with 1 ×10^11^ particles PKH67-labeled circRNA^OVA^-DCpep@EVs (green). LNs were collected 24 h post-injection for cryosectioning and immunostaining against CD11b (red) and DAPI (blue). Confocal microscopy images show overview and magnified views of three representative sites (scale bars: 200 and 50 µm). (F, G) In vivo bioluminescence imaging of C57BL/6 mice after subcutaneous injection of 1×10^11^ particles circRNA^NLuc^@EVs or mRNA^NLuc^@EVs (n=3). Representative images show the time-dependent luciferase expression at the indicated time points (24, 48, 72, 120, and 168 h). Dendritic cell activation was evaluated 24 hours after subcutaneous immunization of mice with the indicated formulations. Single-cell suspensions from inguinal lymph nodes were analyzed by flow cytometry for the surface expression of (H, I) co-stimulatory molecules (CD40, CD86) and (J, K) antigen presentation (MHC I-OVA^+^) on CD11c^+^ cells. (L) C57BL/6 mice (n=5) were immunized subcutaneously on days 0, 5, and 10 with the indicated formulations of circRNA^OVA^-DCpep@EVs (1×10^10^ or 1×10^11^ particles). Splenocytes were isolated on day 20 and restimulated with OVA_257-264_ peptide for 24 h. Intracellular expression of IFN-γ (M) and TNF-α (N) in CD8^+^ T cells was quantified by flow cytometry. (O) Frequency of antigen-specific CD8^+^ T cells was determined using OVA tetramer staining. (P) Cytotoxic activity of splenocytes against B16-OVA target cells was assessed by CCK-8 assay. Data are presented as mean ± SD. Statistical analysis in I, J, M, N, O and P were using one-way ANOVA with a Tukey multiple comparisons test. Statistical analysis in D and G were using two-way ANOVA with a Tukey multiple comparisons test.

To gain deeper insights into the global transcriptional changes, we performed RNA sequencing (RNA-seq) analysis on BMDCs treated with the different vaccine formulations. Differential gene expression analysis identified 1,611 significantly differentially expressed genes in the circRNA^OVA^-DCpep@EVs group versus the control, compared to 574 genes in the circRNA^OVA^@EVs group (**fig. S6A**). A direct comparison between the targeted and non-targeted EV vaccines revealed 185 upregulated and 52 downregulated genes (**fig. S6B**). Gene Ontology (GO) enrichment analysis indicated that the upregulated genes were predominantly involved in innate immune response, inflammatory response, and immune system processes (**fig. S6C**). Consistent with this, KEGG pathway analysis demonstrated that circRNA^OVA^-DCpep@EVs treatment significantly modulated key immune-related pathways, including the TNF signaling pathway and the Toll-like receptor signaling pathway (**fig. S6D**).

An ideal vaccine delivery system must efficiently transport its payload to lymph nodes (LNs), where DCs process and present antigens to prime T cell immunity. To evaluate in vivo LN uptake, PKH67-labeled EVs were administered subcutaneously, and draining LNs were harvested 24 hours post-injection for immunofluorescence analysis. The results demonstrated significantly enhanced co-localization of CD11c^+^ DCs with EVs in LNs from mice treated with circRNA^OVA^-DCpep@EVs compared to PBS controls. Notably, this co-localization was more pronounced in circRNA^OVA^-DCpep@EVs compared to circRNA^OVA^@EVs group (**Fig. 4E**). Consistent with these findings, in vivo imaging system (IVIS) analysis of LNs revealed a stronger fluorescent signal following subcutaneous injection of circRNA^OVA^-DCpep@EVs relative to the circRNA^OVA^@EVs group, although the signal was mainly distributed in the liver (**fig. S7A-B**). To further assess the protein expression mediated by EV-delivered circRNA, we packaged either mRNA^Nluc^ or circRNA^Nluc^ into EVs and monitored nanoluciferase expression in mice over time. Animals injected with circRNA^Nluc^@EVs sustained protein production for over 7 days (**Fig. 4F**). Importantly, in contrast to mRNA, circRNA-driven protein levels continued to increase over the first 72 hours and maintained higher relative fluorescence intensity at 168 hours (**Fig. 4G**).

To evaluate the immunogenicity of EV-delivered circRNA vaccines, C57BL/6 mice were subcutaneously immunized with PBS, EVs, circRNA^OVA^@EVs and circRNA^OVA^-DCpep@EVs. Draining LNs were harvested to assess DC activation status. Flow cytometric analysis revealed that mice immunized with circRNA^OVA^-DCpep@EVs exhibited significant upregulation of the co-stimulatory molecules CD40 and CD86 on CD11c⁺ cells compared to control groups. Notably, the proportion of activated CD11c⁺CD40⁺CD86⁺ DCs was substantially higher in the circRNA^OVA^-DCpep@EVs group than in the circRNA^OVA^@EVs group (**Fig. 4H-I**). Furthermore, circRNA^OVA^-DCpep@EVs immunization resulted in a significantly increased frequency of CD11c⁺MHCI-OVA⁺ cells, indicating enhanced antigen presentation (**Fig. 4J-K**). These findings demonstrate that DCpep modification on the EV membrane significantly promotes DC uptake and antigen presentation capabilities. Subsequent investigations were conducted to characterize the antigen-specific immune response elicited by this vaccine platform. C57BL/6 mice were immunized subcutaneously with circRNA^OVA^-DCpep@EVs on a schedule of three doses administered at 5-day intervals. On day 20 post-initial immunization, mice were euthanized and splenocytes were isolated for ex vivo restimulation with OVA_257-264_ (SIINFEKL) peptide (**Fig. 4L**). Intracellular cytokine staining revealed that the frequencies of IFN-γ⁺ and TNF-α⁺ cells among CD8⁺ T lymphocytes were significantly elevated in the circRNA^OVA^-DCpep@EVs group compared to all other experimental groups (**Fig. 4M-N**). Furthermore, OVA tetramer staining confirmed the induction of a substantially larger population of antigen-specific CD8⁺ T cells in these mice (**Fig. 4O**). To assess functional cytotoxicity, splenocytes from immunized mice were co-cultured with B16-OVA target cells. As summarized in **Fig. 4P**, splenocytes from the circRNA^OVA^-DCpep@EVs group exhibited potent and specific killing activity against B16-OVA cells. Collectively, these findings demonstrate that the circRNA^OVA^-DCpep@EVs vaccine facilitates efficient DC uptake and activation in vivo, leading to the generation of a robust and functional antigen-specific CD8⁺ T cell response.

Collectively, these in vitro and in vivo results demonstrate that the engineered circRNA^OVA^-DCpep@EVs platform facilitates cytosolic antigen delivery, sustains antigen presentation, and induces robust and multifaceted DC activation, thereby eliciting a potent immune response.

### EV-mediated circRNA vaccine elicits prophylactic and therapeutic antitumor immunity with systemic and tumor-infiltrating T cell responses

Melanoma, a prevalent malignancy often characterized as an immunologically “hot tumor” due to its substantial immune cell infiltration, has been a major focus of immunotherapeutic strategies aimed at augmenting T cell activity and increasing intratumoral infiltration of T cells(*20, 21*). To evaluate the prophylactic potential of the circRNA^OVA^-DCpep@EVs vaccine, we established a B16-OVA melanoma model in C57BL/6J mice (**Fig. 5A**). Assessment of tumor growth kinetics and final tumor weights revealed that the circRNA^OVA^-DCpep@EVs group exhibited the most potent suppression of tumor progression compared to both circRNA^OVA^@EVs and the control groups (**Fig. 5B-C**). Notably, one mouse in the circRNA^OVA^-DCpep@EVs group remained largely tumor-free until day 17 (**Fig. 5D**). Immune profiling of splenocytes demonstrated that vaccination with circRNA^OVA^-DCpep@EVs elicited a superior antigen-specific CD8^+^ T cell response (**fig. S8A**). This was evidenced by increased polyfunctional cytokine production (IFN-γ, IL-2, TNF-α) and enhanced granzyme B (GZMB) secretion compared to controls (**Figure 5E and fig. S8B**). ELISpot analysis further confirmed the induction of a more robust antigen-specific T lymphocyte population (**Fig. 5F-G**). Crucially, the circRNA^OVA^-DCpep@EVs vaccine significantly expanded both CD8^+^ central memory (TCM) and effector memory (TEM) T cell subsets in the spleen, providing a mechanistic basis for its sustained prophylactic efficacy (**Fig. 5H-J and fig. S9**).

**Fig. 5.**
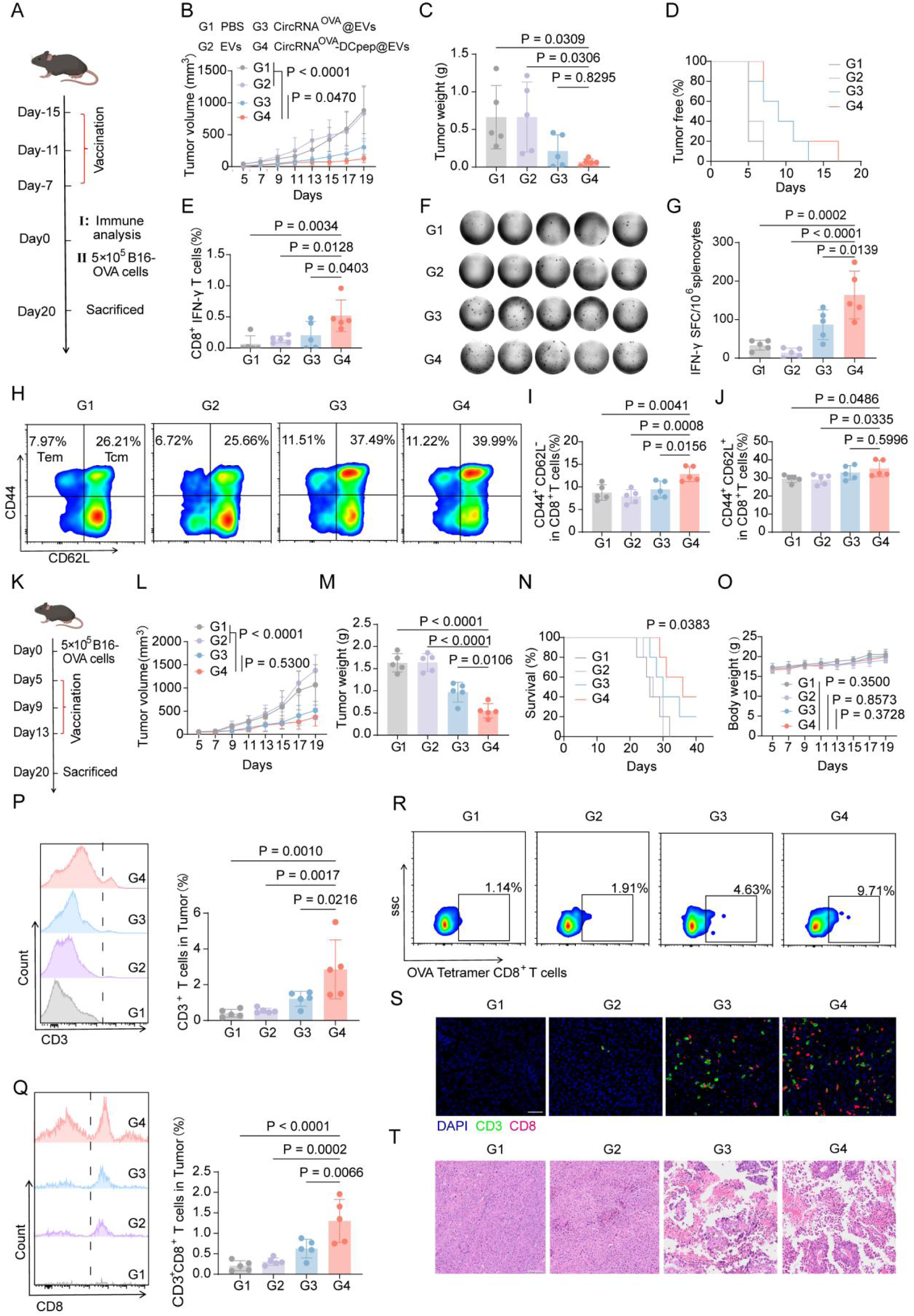
Prophylactic and therapeutic efficacy of the EV-delivered circRNA vaccine in a B16-OVA murine melanoma model. (A) Timeline illustrating the prophylactic immunization regimen and subsequent tumor challenge in the B16-OVA model (n=5 mice per group). C57BL/6 mice were immunized subcutaneously on days -15, -11, and -7 with the indicated formulations. Splenocytes were isolated on day 0 and the remaining mice were challenged with B16-OVA cells, then mice were sacrificed at day 20 and tumor tissue was harvested. (B) Kinetic profiles of tumor growth (n=5 mice per group) and (C) final tumor weights across the different cohorts. (D) Survival curves depicting tumor-free incidence (n=5 mice per group). Flow cytometric evaluation of splenic T cell populations shows (E) the proportion of CD8^+^ IFN-γ^+^ T cells. (F-G) Antigen-specific IFN-γ ELISpot assay using splenic lymphocytes isolated from mice on day 20. (H) Flow cytometric evaluation of splenic T cell populations shows the proportion of (I) CD8^+^ CD44^+^ CD62L^-^ T cells (TEM), and (J) CD8^+^ CD44^+^ CD62L^+^ T cells (TCM) in each treatment group. (K) Timeline of therapeutic immunization regimen and subsequent vaccination in B16-OVA model (n=5 mice per group). C57BL/6 mice were challenged with B16-OVA cells at day 0, and then immunized subcutaneously on days 5, 9, and 13 with the indicated formulations. The mice were sacrificed at day 20 and tumor tissue was harvested. (L) Growth kinetics of the average volumes of B16-OVA tumors in the indicated groups (n=5 mice per group). (M) Tumor weight of different treatment groups. (N) Survival curves of mice in various treatment groups (n=5 mice per group). (O) Body weight curves of mice in various treatment groups (n=5 mice per group). Flow cytometric assessment of T cell populations in tumors: (P) CD3^+^ T cells, (Q) CD8^+^ T cells, and (R) the antigen-specific CD8^+^ population (identified by OVA tetramer staining) in each treatment group. (S) Representative immunofluorescence micrographs of B16F10 tumor sections harvested on day 20. Sections were stained with antibodies against CD3 (green) and CD8 (red), with nuclei counterstained using DAPI (blue). Scale bar: 50 μm. (T) H&E staining of B16F10 tumour tissues. Scale bar, 100 μm. Data are presented as mean ± SD. Statistical analysis in d, g and h were using one-way ANOVA with a Tukey multiple comparisons test. Statistical analysis in c and f were using two-way ANOVA with a Tukey multiple comparisons test. Data are presented as mean ± SD. Statistical analysis in C, E, G, I, J, M, P and Q were using one-way ANOVA with a Tukey multiple comparisons test. Statistical analysis in B, L, N and O were using two-way ANOVA with a Tukey multiple comparisons test.

Based on the promising prophylactic efficacy of circRNA vaccines, we next evaluated their therapeutic potential against established tumors in vivo. A subcutaneous B16-OVA melanoma model was established in C57BL/6 mice, followed by three subcutaneous administrations of the circRNA vaccine post-tumor inoculation. Tumor growth was monitored until day 20, at which point all mice were humanely euthanized (**Fig. 5K**). Analysis of tumor growth kinetics and final tumor weights revealed that both the circRNA^OVA^@EVs and circRNA^OVA^-DCpep@EVs groups exhibited significant suppression of tumor progression (**Fig. 5L-M**) and extended survival (**Fig. 5N**) compared to the PBS and EV-only control groups. Notably, administration of the circRNA^OVA^-DCpep@EVs vaccine did not induce significant body weight loss, indicating a favorable safety profile (**Fig. 5O**).

Given that antigen presentation by APCs is crucial for priming T cell-mediated anti-tumor immunity, we investigated vaccine-induced T cell responses within the tumor microenvironment. Flow cytometric analysis of tumor-infiltrating lymphocytes revealed significantly enhanced infiltration of both CD3^+^ and CD8^+^ T cells in the circRNA^OVA^-DCpep@EVs group compared to other experimental groups (**Fig. 5P-Q)**. To assess antigen-specificity, we quantified cytotoxic T lymphocytes (CTLs) using OVA tetramer staining, which demonstrated a robust induction of tumor-antigen-specific CD8^+^ T cells in the circRNA^OVA^-DCpep@EVs group (**Fig. 5R and fig. S10**). Immunofluorescence staining of tumor sections corroborated these findings, showing a substantial increase in CD3⁺CD8⁺ T cell density relative to both PBS and EVs control groups (**Fig. 5S)**. H&E staining revealed extensive tumor cell necrosis in the circRNA^OVA^-DCpep@EVs treatment group, characterized by nuclear pyknosis and chromatin condensation (**Fig. 5T)**.

Collectively, these data demonstrate that EV-mediated delivery of circRNA-encoded antigens elicits potent antitumor immunity. The DCpep engineering strategy significantly enhances this effect, promoting the generation of systemic immune memory for tumor prevention and driving robust T cell infiltration and tumor destruction within the established tumor microenvironment.

### EV-mediated delivery of BNP-encoding circRNA mitigates doxorubicin-induced myocardial fibrosis

Exogenous B-type natriuretic peptide (BNP) exerts therapeutic effects in heart failure by promoting vasodilation, reducing cardiac load, inducing natriuresis, and inhibiting renin-angiotensin-aldosterone system (RAAS) activation, thereby alleviating hemodynamic dysfunction and pathological cardiac remodeling(*22*). Clinically, BNP supplementation rapidly improves symptoms and reduces mortality in patients with chronic heart failure (CHF)(*23*). Therefore, we utilized an EV-based circRNA delivery platform to overexpress human BNP (hBNP) in a murine model of doxorubicin (DOX)-induced cardiotoxicity. Male C57BL/6J mice were treated according to the regimen outlined in **Fig. 6A**. Prior to euthanasia, cardiac function was assessed via echocardiography.

**Fig. 6.**
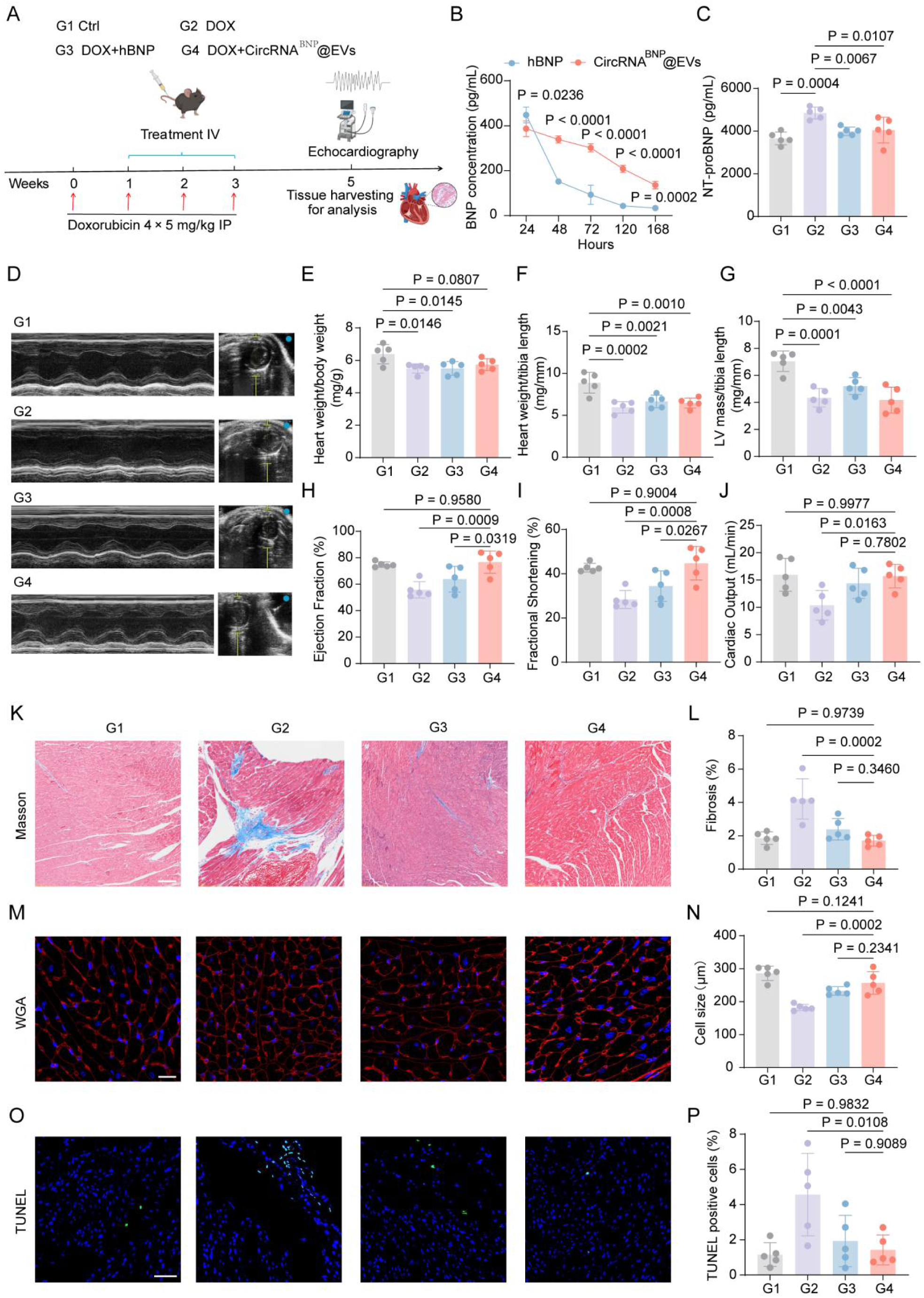
Systemic administration of BNP attenuates doxorubicin-induced cardiotoxicity in mice. (A) Timeline of the Doxorubicin-Induced Heart Failure Model and Therapeutic Interventions. C57BL/6 mice were randomly assigned to four treatment groups, receiving either PBS or DOX (5 mg/kg, i.p., four doses). Corresponding therapeutic interventions: PBS, hBNP, or circRNA^BNP^@EVs were administered intravenously in three doses. Echocardiography analysis was performed prior to euthanasia. The overall experimental design is summarized in the schematic timeline. (B) Serum levels of hBNP measured by ELISA at indicated time points. (C) Serum NT-proBNP levels quantified by ELISA two weeks post-treatment. (D) Representative two-dimensional M-mode echocardiographic images of left ventricle. (E) Heart weight-to-body weight ratio. (F) Heart weight-to-tibia length ratio. (G) Left ventricular (LV) mass normalized to tibia length. (H) Ejection fraction (%) calculated from echocardiography. (I) Fractional shortening (%) from echocardiography. (J) Cardiac output derived from echocardiographic measurements. (K) Representative Masson’s trichrome staining of myocardial sections (scale bar = 100 μm). (L) Quantification of myocardial fibrosis area (%) from (K) (n = 5 per group). (M) Representative wheat germ agglutinin (WGA)-stained sections for cardiomyocyte cross-sectional area (scale bar = 25 μm). (N) Quantification of cardiomyocyte size from (M) (n = 5 per group). (O) Representative TUNEL-stained myocardial sections (scale bar = 50 μm; black arrowheads indicate TUNEL-positive nuclei). (P) Quantitative analysis of TUNEL-positive nuclei per field from (O) (n = 5 per group). Data are presented as mean ± SD. Statistical analysis in C, E, F, G, H, I, J, L,N and P were using one-way ANOVA with a Tukey multiple comparisons test. Statistical analysis in B was using two-way ANOVA with a Tukey multiple comparisons test.

We first evaluated the pharmacokinetics of systemically delivered hBNP. Serum levels of hBNP were monitored after tail vein injection of recombinant hBNP protein or circRNA^BNP^@EVs. As shown in **Fig. 6B**, EV-mediated delivery of circRNA^BNP^ led to sustained hBNP expression detectable for up to one week, contrasting with the transient presence observed with direct protein administration. In DOX-treated mice, circulating NT-proBNP levels were significantly reduced (**Fig. 6C**).

Echocardiographic and morphometric analyses revealed that DOX challenge induced progressive weight loss and cardiac atrophy (**Fig. 6D-G**). Notably, mice receiving combined DOX and circRNA^BNP^@EVs exhibited improved cardiac function compared to DOX-only controls, as evidenced by increased ejection fraction, fractional shortening, and cardiac output. This functional recovery surpassed that achieved by hBNP protein injection (**Fig. 6H-J**). Histopathological assessment further supported the cardioprotective effect of sustained BNP expression. The circRNA^BNP^@EVs group showed attenuated myocardial fibrosis (**Fig. 6K-L**), increased cardiomyocyte cross-sectional area (**Fig. 6M-N**), and a decreased frequency of TUNEL-positive nuclei (**Fig. 6O-P**). Together, these findings demonstrate that EV-facilitated circRNA delivery enables prolonged BNP expression and effectively alleviates DOX-induced cardiac injury and structural remodeling.

## Discussion

Efficient endogenous loading of circRNA into EVs requires the coordinated enhancement of both intracellular transcript abundance and selective enrichment within vesicles. Although circRNA has emerged as a promising modality for RNA therapeutics owing to its enhanced stability and capacity for sustained protein expression, its broader application has been constrained by two fundamental limitations: inefficient intracellular expression and poor delivery efficiency. Existing circRNA expression systems, including Alu-mediated back-splicing and intron-based circularization strategies, often generate low levels of translatable circRNA and substantial linear byproducts(*24, 25*). While the Tornado system represents a significant advance by leveraging tRNA splicing machinery, its circularization efficiency and overall expression output remain substantially lower than those of linear mRNA, thereby limiting its utility for functional delivery(*10, 11*).

In this study, we address these limitations through an integrated strategy that simultaneously enhances circRNA expression and EV-mediated delivery. By optimizing intracellular circularization and translation, we substantially improve circRNA output, overcoming a key bottleneck associated with endogenous processing systems. In parallel, incorporation of an RNA-binding protein–mediated sorting mechanism, together with enhanced EV biogenesis, enables efficient encapsulation of circRNA without compromising vesicle integrity. The requirement for all components in achieving functional delivery highlights the importance of coordinated control over RNA production, vesicle loading, and EV secretion, rather than optimizing each step in isolation.

Importantly, the two application scenarios explored in this study further illustrate the functional breadth of this platform. In the context of cancer immunotherapy, EV-mediated delivery of circRNA vaccines enables efficient antigen expression and cross-presentation in dendritic cells, leading to robust activation of antigen-specific T cell responses and effective tumor control in both prophylactic and therapeutic settings. In parallel, delivery of BNP-encoding circRNA demonstrates the applicability of this system for protein replacement–based therapies, where sustained expression is required to achieve therapeutic benefit.

Notably, these applications involve circRNA cargos with distinct lengths and biological functions, underscoring the versatility of this platform. The ability to support both immune activation and tissue-protective effects suggests that coordinated enhancement of circRNA expression and EV loading can extend circRNA therapeutics across fundamentally different disease contexts. More broadly, the capacity to efficiently deliver relatively large coding circRNAs addresses a key limitation associated with EV-mediated RNA delivery.

Compared to conventional RNA delivery systems, including lipid nanoparticles, EV-based delivery offers intrinsic advantages in biocompatibility and cellular uptake. Our findings extend these advantages by enabling endogenous loading through coordinated regulation of circRNA biogenesis and vesicle sorting, thereby overcoming the limitations of exogenous loading approaches. Conceptually, this work shifts the paradigm from passive cargo loading toward programmable cargo biogenesis and sorting, providing a new framework for RNA delivery.

In summary, we establish an EV-based platform that enables efficient circRNA delivery through in situ biogenesis and RNA-guided sorting. By overcoming key barriers in circRNA expression and loading, this work provides a generalizable strategy for durable RNA therapeutics and expands the potential applications of circRNA across diverse therapeutic modalities.

## Materials and Methods Plasmid design

All DNA fragments were chemically synthesized (Tsingke Bio, China) with codon optimization for mammalian cell expression, employing established protocols(*26*). Each gene was subsequently cloned into a pcDNA3 expression vector under the control of the CMV promoter. The circRNA expression plasmid was constructed by cloning a synthetic sequence comprising the 5’ UTR, IRES, Kozak sequence, the CDS, C/D box motif, and the 3’ UTR, flanked by twister ribozyme sequences. Subsequently, spacer sequences of varying lengths were introduced between the CMV promoter and the twister ribozyme structural elements. To improve the translation efficiency of circRNA expression vectors, a WPRE element was incorporated downstream of the 3’ twister ribozyme sequence, while LTR sequences were positioned flanking the entire expression cassette.

The following additional key plasmids were also utilized: pcDNA3-CD63-Snu13 expresses a fusion protein consisting of the exosomal marker CD63 with an N-terminal HA tag and the RNA-binding protein Snu13(*27*) conjugated to its C-terminus. pcDNA3-CD63-Nluc encodes the nanoluciferase reporter gene conjugated to the C-terminus of CD63. pcDNA3-DCpep-LAMP2B contains an engineered LAMP2B construct featuring N-terminal Flag tag, a glycosylation signal and DCpep peptide(*28*). Additionally, an exosome production enhancer plasmid was used to express a fusion protein incorporating STEAP3, SDC4, and NadB(*18*).

### Cell culture and Plasmid DNA transfection

HEK-293T cells were cultured in DMEM supplemented with 10% fetal bovine serum (FBS) and 1% penicillin/streptomycin. DC2.4 cells, BMDC cells and B16F10-OVA cells were cultured in RPMI 1640 medium containing 10% FBS and 1% penicillin/streptomycin. All cell lines were incubated at 37 ℃ under 5% CO2.

For transfection experiments, HEK-293T cells were plated in 6-well plates at a density of 60-70% confluency prior to transfection. A DNA-polyethyleneimine (PEI) complex was prepared by combining 2 μg of total DNA with 5 μL of PEI, followed by brief vortexing and a 30 minute incubation at room temperature. A volume of 200 μL of this complex was applied per well. The procedure was proportionally scaled when using alternative culture vessels such as 12-well plates, 6-well plates, 10-cm dishes, or 15-cm dishes. Before transfection, the existing cell culture medium was replaced with fresh DMEM supplemented with 10% FBS. The cells were then incubated with the transfection mixture for 8 to 16 hours at 37 ℃ under 5% CO2. Following this incubation, the medium was replaced again with fresh pre-warmed medium to support subsequent expression of the gene of interest.

### RNA isolation and RT-qPCR

RNA was isolated from tissues or cultured cells using FreeZol Reagent (Vazyme, China) following the manufacturer’s protocol. RNA concentration and purity were determined by measuring absorbance at 260 nm and 280 nm using a spectrophotometer (NanoDrop Technologies, USA). cDNA synthesis was carried out using the Hifair® III 1st Strand cDNA Synthesis Kit (Yeasen, China). Quantitative real-time PCR was performed with Hieff® qPCR SYBR Green Master Mix (Yeasen, China) on a QuantStudio 5 Detection System (Applied Biosystems, USA). GAPDH expression was used as an endogenous control to normalize the expression levels of target mRNAs or circRNAs. The primer sequences were shown in **Table S3**.

To confirm the circular nature and resistance to exonuclease degradation of the circRNA molecules, total RNA (2 μg) was digested with RNase R (5 U/μg; Novoprotein, China) at 37°C for 15 minutes. The resulting RNA was subsequently subjected to RT-qPCR analysis to evaluate its stability and structural characteristics.

### Absolute qPCR

Absolute qPCR was performed to determine the copy number of circRNA in both circRNA-overexpressing HEK-293T cells and EVs. Briefly, a 399-bp fragment spanning the back-splice junction of the circRNA was cloned into the pCDNA3.1 vector. Serial tenfold dilutions of the plasmid were used to generate a standard curve by real-time PCR. The copy number of the standard was calculated using the formula: copy number (copies/μL) = [6.02 × 10²³ × plasmid concentration (ng/μL) × 10⁻⁹] / [(vector molecular weight + insert molecular weight) × 660]. The standard curve was constructed by plotting the logarithm of the initial template copy number (X-axis) against the threshold cycle (Ct) value (Y-axis), yielding a linear equation: Y = aX + b. The Ct values obtained from circRNA amplifications were then converted into absolute copy numbers based on this standard curve.

### RNA immunoprecipitation (RIP)

RIP assays were carried out using the Magna RIP® RNA-Binding Protein Immunoprecipitation Kit (Merck Millipore, USA) in accordance with the manufacturer’s protocol. HEK-293T cells were cultured to an appropriate density in 15-cm dishes and co-transfected with plasmids pcDNA3-circRNA^OVA^-C/D box and pcDNA3-CD63-Snu13 at a 1:1 ratio. Following transfection, cells were lysed in 100 μL of RIP lysis buffer supplemented with RNase inhibitors. The resulting lysates were subjected to immunoprecipitation by incubation with magnetic beads conjugated either with anti-HA tag antibody or normal IgG (as a negative control) on a rotary shaker overnight at 4 °C. RNAs co-precipitated with the target complexes were subsequently isolated and analyzed by RT-qPCR and agarose gel electrophoresis.

### CircRNA transfer assay

To investigate exosome-mediated RNA transfer, HEK-293T cells were seeded in 6-well plates and co-transfected with a plasmid mixture comprising 2 μg of pcDNA3-circRNA^Nluc^-C/D box, 1 μg of pcDNA3-CD63-Snu13, and 1 μg of pcDNA3-Enhancer, using 10 μL of PEI transfection reagent according to the manufacturer’s instructions. The culture supernatant was harvested from the producer cells 48 hours after medium replacement post-transfection.

Following centrifugation to remove cellular debris, the clarified supernatant was applied to recipient cells plated in 24-well plates at 60-80% confluency. After 24 hours of incubation, Nano Luciferase activity in the recipient cells was assessed using the Nano Luciferase Reporter Gene Assay Kit (Biolight, China). Briefly, 200 µL of cell suspension was mixed with 200 µL of quantification reagent, prepared by diluting 4 µL of Nano Luciferase Assay Substrate in 200 µL of Nano Luciferase Assay Buffer. The mixture was homogenized by pipetting, and 100 µL of the resulting solution was transferred to a white 96-well plate. Luminescence was measured at 25 °C using a Tecan Infinite E Plex system with an integration time of 1 second per well. Results are expressed as relative light units (RLU).

### The preparation of EVs

EVs were isolated from HEK-293T cells cultured in 15-cm dishes. Briefly, the cells were transfected with a combination of plasmids (9.5 μg of pcDNA3-circRNA^OVA^-C/D box, 4.75 μg of pcDNA3-CD63-Snu13, 2 μg of pcDNA3-DCpep-LAMP2B, and 4.75 μg of pcDNA3-Enhancer) using 50 μL of PEI transfection reagent in accordance with the manufacturer’s protocol.

Following a 24-hour post-transfection interval, the culture medium was replaced with fresh DMEM. The conditioned supernatant was subsequently harvested 48 hours after this medium change for EVs purification. EVs derived from the supernatant fluids of HEK-293T cells were isolated via differential centrifugation. As a control, mock EVs were also obtained from untransfected HEK-293T cells. The supernatant was subjected to a series of centrifugation steps: initially at 300 × g for 10 min, followed by 2000 × g for 30 min, and then 10,000 × g for 60 min, all performed at 4 °C to eliminate cells and cellular debris. The clarified supernatant was subsequently filtered through a 0.22-μm membrane filter (Millipore, USA). Ultracentrifugation of the filtrate was carried out at 200,000 × g for 120 min at 4 °C using an Optima MAX-XP ultracentrifuge (Beckman Coulter, USA). The final EV pellet was resuspended in phosphate-buffered saline (PBS) for subsequent applications.

### Characterization of EVs

The characterization of purified EVs was performed using established biomarkers via western blot analysis as previous descried(*29, 30*). The following primary antibodies were applied: anti-CD63 (1:1000, ab134045, Abcam), anti-CD9 (1:1000, ab263019, Abcam), anti-TSG101 (1:1000, ab125011, Abcam), anti-Hsp70 (1:1000, ab181606, Abcam), anti-HA tag (1:5000, 51064-2-AP, Proteintech), and anti-Flag tag (1:5000, 66008-4-Ig, Proteintech). Additionally, the functional validation of the encapsulated circRNA’s protein expression capability was assessed by western blot analysis. Following transfection into HEK-293T cells, protein expression from the EV-delivered circRNA was detected using an anti-His tag antibody (1:5000, 66005-1-Ig, Proteintech). The morphological features of EVs were examined using transmission electron microscopy (TEM, Talos L120C, ThermoFisher, USA). Briefly, purified EVs were fixed with 4% paraformaldehyde (PFA). A 10 µL aliquot of the EV suspension was applied to a copper grid and incubated at room temperature for 10 minutes. After washing three times with sterile distilled water, the grid was negatively stained with 10 µL of 2% uranyl acetate solution for 1 minute.

Excess liquid was carefully removed using filter paper, and the grid was air-dried under an incandescent lamp for 2 minutes. Imaging was performed at an accelerating voltage of 80 kV. The size distribution and particle concentration of EVs were determined by nanoparticle tracking analysis (NTA, NanoSight NS300, Malvern, UK). The protein concentration of purified EVs was quantified using a Pierce BCA Protein Assay Kit (Beyotime, China) in strict accordance with the manufacturer’s instructions.

### RNA Protection Assay

CircRNA^OVA^-DCpep@EVs (1 × 10^11^ particles) were treated with 20 µL of RNase A/T1 mixture at 37 °C for 30 minutes, either in the presence or absence of 1% Triton X-100. Following enzyme inactivation at 75 °C for 5 minutes, RNA was immediately extracted using the exosome RNA Purification Kit (EZBioscience, USA). An equal volume of purified RNA was reverse transcribed as described previously. The absolute copy number of circRNA was subsequently quantified via absolute qPCR.

### In vitro evaluation of DCs uptake, antigen presentation, and activation

BMDCs were isolated and cultured using a standard protocol. Briefly, female C57BL/6 mice were euthanized and sterilized with 70% ethanol. The femurs and tibias were dissected to harvest bone marrow cells. After erythrocyte lysis using red blood cell lysis buffer, the remaining cells were resuspended in RPMI-1640 complete medium supplemented with 20 ng/mL granulocyte-macrophage colony-stimulating factor (GM-CSF). On day 3 of culture, an equal volume of fresh medium containing 20 ng/mL GM-CSF was added. On day 6, half of the culture medium was replaced with fresh GM-CSF-containing medium. BMDCs were harvested and seeded into plates between days 7 and 9 for subsequent experiments.

To evaluate cellular uptake, circRNA^OVA^-DCpep@EVs were fluorescently labeled using the PKH-67 Green Fluorescent Cell Linker Kit (Beyotime, China) according to the manufacturer’s instructions. A pellet containing 1 × 10^9^ PKH67-labeled EVs was resuspended and co-incubated with BMDCs seeded in 24-well plates at 37 °C for 0, 2, 4, and 8 hours. Following incubation, cells were washed with PBS and fixed with 4% paraformaldehyde. Nuclei were counterstained with DAPI, and EV internalization was visualized using a confocal microscope (LSM900, Zeiss, Germany).

For antigen presentation analysis, BMDCs were seeded in 24-well plates and treated with 5×10^9^ particles of either circRNA^OVA^@EVs or circRNA^OVA^-DCpep@EVs. At time points of 0, 6, 12, 24, 36, and 48 hours post-treatment, the expression of the MHCI-OVA complex was analyzed by flow cytometry using T-Select H-2Kb OVA Tetramer-SIINFEKL-APC antibody (BetterGen, China).

To assess immune activation, BMDCs were seeded in 24-well plates and stimulated with 5 × 10^9^ particles of the respective EV preparations. After 24 hours, cells were collected and stained with the following fluorescently conjugated antibodies: FITC anti-mouse CD11c (557400, BD Biosciences), PE-Cy7 anti-mouse CD86 (560582, BD Biosciences), BV421 anti-mouse CD40 (562846, BD Biosciences), and PE anti-mouse MHCⅠ (557000, BD Biosciences). After incubation for 30 minutes at room temperature, cells were washed and analyzed using a NovoCyte Quanteon flow cytometer (Agilent, China). In parallel, total RNA was extracted from harvested cells using FreeZol Reagent. The expression of cytokines (IFN-γ, IL-6, and TNF-α) was detected via RT-qPCR. Additionally, cell culture supernatants were centrifuged to remove debris, and cytokine levels were quantified using the Bio-Plex Cytokine Assay Kit (Bio-Rad, USA). To analyze differential gene expression in BMDCs before and after activation, RNA-seq was performed. RNA integrity was verified, and libraries were prepared and sequenced on an Illumina platform. Bioinformatic analysis was conducted to identify significantly differentially expressed genes, with functional enrichment analysis performed to elucidate involved biological pathways.

### In vivo evaluation of draining lymph node uptake, antigen presentation, and activation

To evaluate the in vivo uptake of circRNA^OVA^-DCpep@EVs by inguinal lymph nodes, C57BL/6 mice received a subcutaneous injection of 1 × 10^11^ PKH67-labeled EVs into the dorsal region. After 24 hours, the LNs were harvested, embedded in optimal cutting temperature (OCT) compound, and snap-frozen. Cryosections of 8 μm thickness were prepared and washed thoroughly with PBS to remove residual OCT. Subsequently, sections were blocked with 5% bovine serum albumin (BSA) and 0.5% Triton X-100 in PBS for 1 hour at room temperature.

Nuclei were counterstained with DAPI, and imaging was performed using a slide scanning system (VS200, Olympus, Japan). Image analysis was conducted with SlideViewer software.

Additionally, fluorescence imaging and quantification of LNs were carried out using an IVIS Spectrum imaging system (PerkinElmer, USA).

To assess antigen presentation and activation of dendritic cells, inguinal LNs were collected 24 hours post-EVs injection. Single-cell suspensions were prepared by mechanically dissociating the nodes through a 70-μm strainer. The resulting cells were stained with the following fluorescently conjugated antibodies: FITC anti-mouse CD11c, PE-Cy7 anti-mouse CD86, BV421 anti-mouse CD40, and H-2Kb OVA Tetramer-SIINFEKL-APC, following standard staining protocols. Finally, flow cytometric analysis was performed to evaluate cell populations and activation states.

### In vivo assessment of protein expression mediated by circRNA@EVs

To validate the in vivo protein expression capability of circRNA encapsulated within extracellular vesicles, C57BL/6 mice were administered a subcutaneous injection of 1 × 10^11^ circRNA^NLuc^@EVs or mRNA^NLuc^@EVs into the dorsal region. At 24, 48, 72, 120, and 168 hours post-injection, the mice received an intraperitoneal injection of 100 μL Fluorofurimazine (Selleck, USA) at a final concentration of 333 nmol(*31*). Bioluminescence imaging was performed 15 minutes after substrate administration using an IVIS Spectrum imaging system.

### Antitumor Immunotherapy

For the therapeutic model of subcutaneous melanoma, C57BL/6 mice were inoculated subcutaneously with 5 × 10^5^ murine B16-OVA melanoma cells. Five days post-inoculation, the mice were randomly assigned to different experimental groups (n = 5) and received the following treatments every four days for a total of three administrations: 100 μL PBS, 1 × 10^11^ naïve EVs, 1 × 10^11^ circRNA^OVA^@EVs, or 1 × 10^11^ circRNA^OVA^-DCpep@EVs. Mice exhibiting severe morbidity were euthanized prior to the endpoint. All remaining mice were sacrificed on day 20 post-tumor inoculation, and tumors and spleens were harvested for further analysis. In the prophylactic vaccination model, C57BL/6 mice were first administered the vaccine subcutaneously (every four days for three doses) and then randomly divided into four groups (n = 5). Seven days after the final vaccination, the mice were challenged subcutaneously with 5 × 10^5^ B16-OVA cells. Any mouse showing signs of severe morbidity was humanely euthanized; all others were sacrificed on day 20 after tumor inoculation. Tumors and spleens were collected for subsequent evaluation. Tumor volume was monitored every two days using a digital caliper and calculated according to the formula: V = (length × width²) / 2, expressed in mm³. Body weight was also recorded regularly throughout the study period.

Spleens were homogenized and passed through a 70-μm cell strainer to obtain single-cell suspensions. Splenic lymphocytes were isolated using a murine lymphocyte separation solution (Dakewe, China). Erythrocytes were lysed by incubation with red blood cell lysis (Biosharp, China) buffer for 5 minutes at 4 °C, followed by washing with RPMI 1640 medium. The purified lymphocytes were resuspended in RPMI 1640 medium supplemented with 10% FBS, 2 mM L-glutamine, and 1% penicillin/streptomycin. For tumor tissue processing, samples were minced into small fragments and digested with 250 μg/mL collagenase IV (MCE, USA) and 120 μg/mL DNase I (MCE, USA) at 37 °C for 1 hour. The digested tissue was then filtered through a 70-μm cell strainer to generate a single-cell suspension for subsequent analysis.

### IFN-γ ELISpot Assay

The IFN-γ ELISpot assay was conducted using Mouse IFN-γ Precoated ELISPOT Kit (Dakewe, China). Briefly, 96-well ELISpot plates were rinsed with PBS and coated with purified rat anti-mouse IFN-γ monoclonal antibody overnight at 4 °C. Mouse splenocytes were seeded into the plates at a density of 5 × 10⁵ cells per well. Cells were then stimulated with an OVA_257-264_ peptide (MCE, USA) at a concentration of 4 mg/mL. Dimethyl sulfoxide (DMSO) and 500 ng/mL PMA + 10 μg/mL Ionomycin were used as negative and positive controls, respectively. After 24 hours of incubation, the plates were incubated with a biotinylated detection antibody, followed by development with alkaline phosphatase-conjugated streptavidin and NBT/BCIP substrate. Spot-forming units (SFUs) were quantified using an automated ELISpot reader (Mabtech IRIS 53, Sweden).

### Intracellular Cytokine Staining (ICS)

ICS assay was performed according to a previously established protocol(*26*). Briefly, mouse splenic lymphocytes were seeded into 96-well plates at a density of 2 × 10^6^ cells per well and stimulated with 4 mg/mL OVA_257-264_ peptide. Dimethyl sulfoxide (DMSO) was used as a negative control, and a combination of 500 ng/mL phorbol 12-myristate 13-acetate (PMA) and 10 μg/mL ionomycin served as a positive control. Cells were incubated for 2 hours at 37 °C. Brefeldin A (BioLegend, USA) was then added, and incubation continued for 16 hours at 37 °C. Thereafter, cells were harvested and stained with the following fluorescently conjugated anti-mouse antibodies for 30 minutes at room temperature in the dark: CD3-FITC (553061, BD Biosciences), CD4-PerCP (550954, BD Biosciences), and CD8-PE-Cy7 (552877, BD Biosciences). Cells were then fixed and permeabilized using Cytofix/Cytoperm reagent (BD Biosciences) for 30 minutes in the dark. Intracellular staining was performed using anti-mouse IFN-γ-APC (554413, BD Biosciences), IL-2-BV421 (562969, BD Biosciences), TNF-α-PE (554419, BD Biosciences) and GZMB-AF700 (372222, Biolegend) for 1 hour at 4 ℃ protected from light. Samples were acquired on a flow cytometer and analyzed using CytExpert software.

To evaluate the formation of memory T cells following vaccination with circRNA^OVA^-DCpep@EVs vaccine, splenic lymphocytes (2 × 10^6^ cells per well) were seeded in 96-well plates and stimulated with 4 mg/mL OVA_257-264_ peptide, using DMSO as a negative control, for 24 hours at 37 ℃. Cells were then harvested and stained with a panel of antibodies including anti-mouse CD3-FITC, CD4-PerCP, CD8-PE-Cy7, CD44-AF700 (560567, BD Biosciences), and CD62L-BV605 (563252, BD Biosciences) for 30 minutes in the dark. After washing, samples were analyzed by flow cytometry.

### In vitro cytotoxicity assay

An in vitro cytotoxicity assay was conducted to evaluate the antitumor efficacy of circRNA^OVA^-DCpep@EVs vaccine formulations. A B16-OVA tumor-bearing mouse model and treatment was administered as mentioned before. The splenocytes were first pulsed overnight with OVA peptide antigen for 24 h and then co-cultured with B16-OVA target cells at an effector-to-target ratio of 10:1 for 24 hours. Following incubation, non-adherent cells were discarded, and adherent cells were gently washed with PBS. Specific cytotoxicity was quantitatively assessed using the CCK-8 assay.

### BNP-based intervention in a doxorubicin-induced cardiotoxicity model

Six-to eight-week-old male C57BL/6 mice were randomly assigned to four experimental groups (n = 5 per group): (1) PBS control, (2) doxorubicin (DOX) only, (3) DOX + recombinant human BNP (hBNP), and (4) DOX + circRNA^BNP^@EVs). To establish a model of chronic cardiotoxicity, mice received intraperitoneal (i.p.) injections of DOX (5 mg/kg) once weekly for four consecutive weeks. Control animals received equivalent volumes of PBS on the same schedule. During weeks 2, 3, and 4 of the DOX regimen, each group was administered the corresponding therapeutic or vehicle via intravenous (i.v.) injection: PBS (100 µL) for groups 1 and 2, hBNP (50 µg in 100 µL PBS) for group 3, and circRNA^BNP^@EVs (1 × 10^11^ particles in 100 µL PBS) for group 4. Cardiac function was assessed by echocardiography at week 6, after which animals were euthanized and heart tissues were collected for histological evaluation of myocardial fibrosis.

### Enzyme-linked immunosorbent assay (ELISA)

Serum levels of human BNP (hBNP) and N-terminal pro-BNP (NT-proBNP) were quantified using commercially available ELISA kits (Elabscience, China). Measurements were performed according to the manufacturer’s protocols. Briefly, samples and standards were added to pre-coated wells, incubated with detection antibodies, and developed with substrate solution.

Absorbance was read at 450 nm using a Tecan Infinite E Plex microplate reader (Tecan Trading AG, Switzerland). Concentrations were determined based on standard curves generated for each assay.

### Histological analysis

Cardiac tissues were processed for histological examination as previously described. Briefly, heart samples were fixed overnight in 4% formalin at room temperature, transferred to phosphate-buffered saline (PBS) at 4°C, and subsequently embedded in paraffin. Thin sections were mounted on glass slides for staining. Paraffin-embedded heart sections were subjected to TUNEL assay and Masson’s trichrome staining, and images were acquired using an Olympus BX43F light microscope. Quantification of myocardial fibrosis was performed on Masson’s trichrome-stained sections using ImageJ software. TUNEL-positive cells were manually enumerated by an observer blinded to experimental groups, and total nuclei counts were obtained via ImageJ-based analysis. High-resolution images of wheat germ agglutinin (WGA)-stained sections were captured with a Zeiss LSM 900 confocal microscope under a 40× objective.

## Statistical Analysis

Statistical analyses were performed using GraphPad Prism software (version 8.0).

Comparisons among more than two groups were conducted using one-way analysis of variance (ANOVA) with a Tukey’s multiple comparisons test, while differences between two groups were assessed using a two-tailed unpaired Student’s t-test. Data are presented as mean ± standard deviation (SD).

## Acknowledgements

This work was supported by the National Natural Science Foundation of China (32271420 and 32322045 to Z. L.), China Postdoctoral Science Foundation (2025M772902 to M. L.) and Doctoral Research Foundation of Dongguan People’s Hospital (K202501 to L. L.) . We thank BioRender.com for the preparation of the figures.

## Author Contributions

M. L., Y.P. and M.C. contributed equally to this work. Project design and supervised by Z.L.; experiment performing by M.L., Y.P., M.C., J.D., F. W., Y.L. and R.Z.; data analysis by M.L. and Y.P.; picture drawing by M.L. and M.C.; Writing by M.L; materials and reagents contributed by M.C., C.S.; review and editing by Z.L. All authors have read and agreed to the final version of this manuscript.

## Data, code, and materials availability

All data are available in the main text or the supplementary materials.

## Supplementary Materials

This PDF file includes: Figs. S1 to S10

Tables S1 to S3

