## Supplementary material for "Extracellular Vesicles Enable CircRNA Delivery via in situ Biogenesis and Sorting": Figs. S1 to S10 and Tables S1 to S3

Minchao Li et al.

**This PDF file includes:**

Figs. S1 to S10
Tables S1 to S3

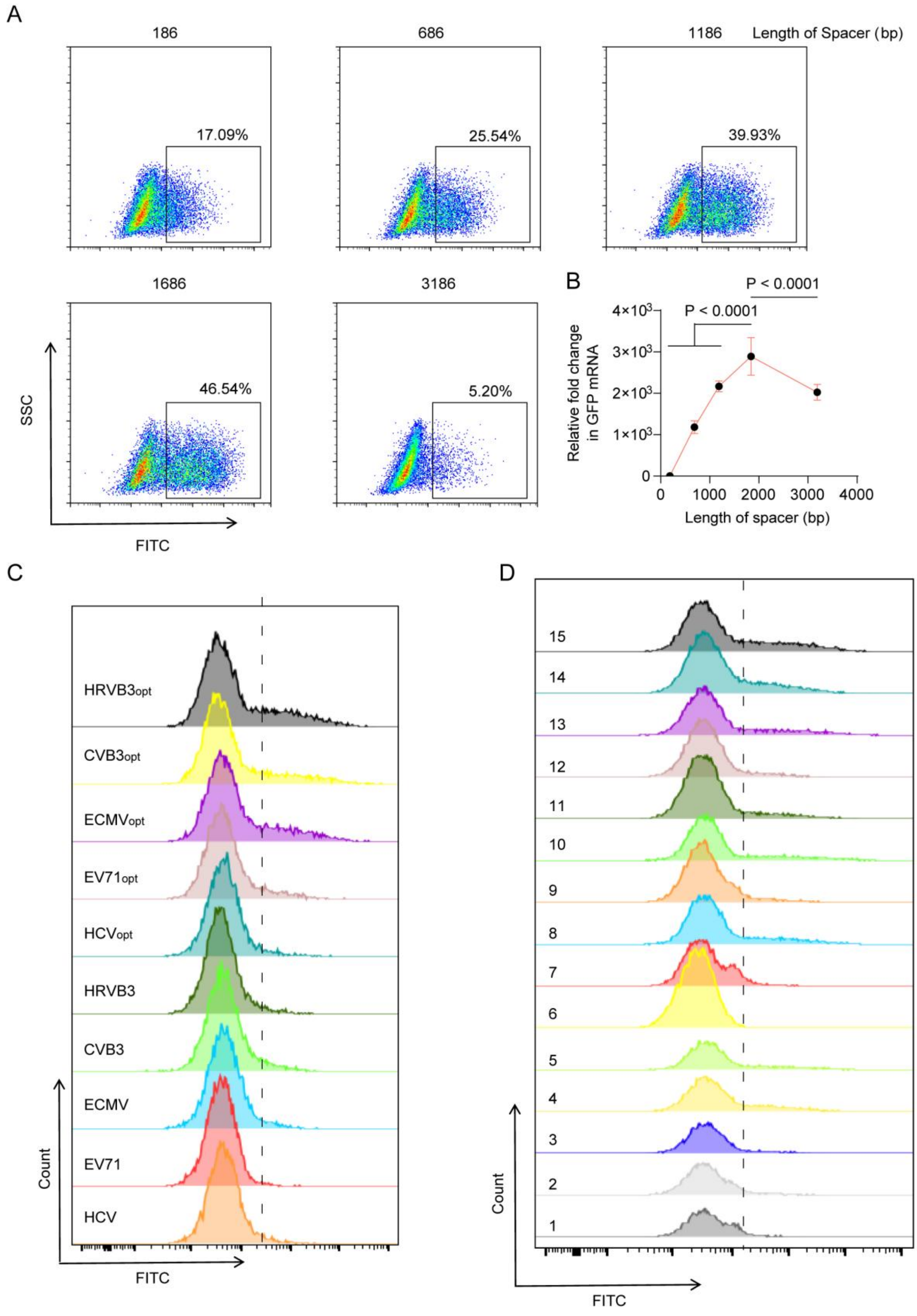


**fig. S1 The optimization of circRNA expression vector.** (A) Length-Dependent Enhancement of GFP Expression by Engineered Spacer Sequences. The HEK-293T cells were treated with different circRNA expression vector insert various length of spacer. The GFP expression efficiency was quantified by flow cytometry. (B) Comparative Analysis of GFP Transcript Levels from Various circRNA Vectors. (C) The performance of diverse IRES sequences within the circRNA expression vector was assessed in HEK-293T cells. Following transfection, GFP intensity was measured quantitatively via flow cytometry. (D) The performance of WPRE and the LTR element within the circRNA expression vector was assessed in HEK-293T cells. The GFP intensity was measured quantitatively via flow cytometry. Three vector sets (#1: plasmids 1-5; #2: plasmids 6-10; #3: plasmids 11-15) were constructed, each containing the same series of IRES elements in the order of HCV, EV71, ECMV, CVB3, and HRV-B3. Data are presented as mean ± SD. Statistical analysis in B was using two-tailed unpaired Student’s t-tes.

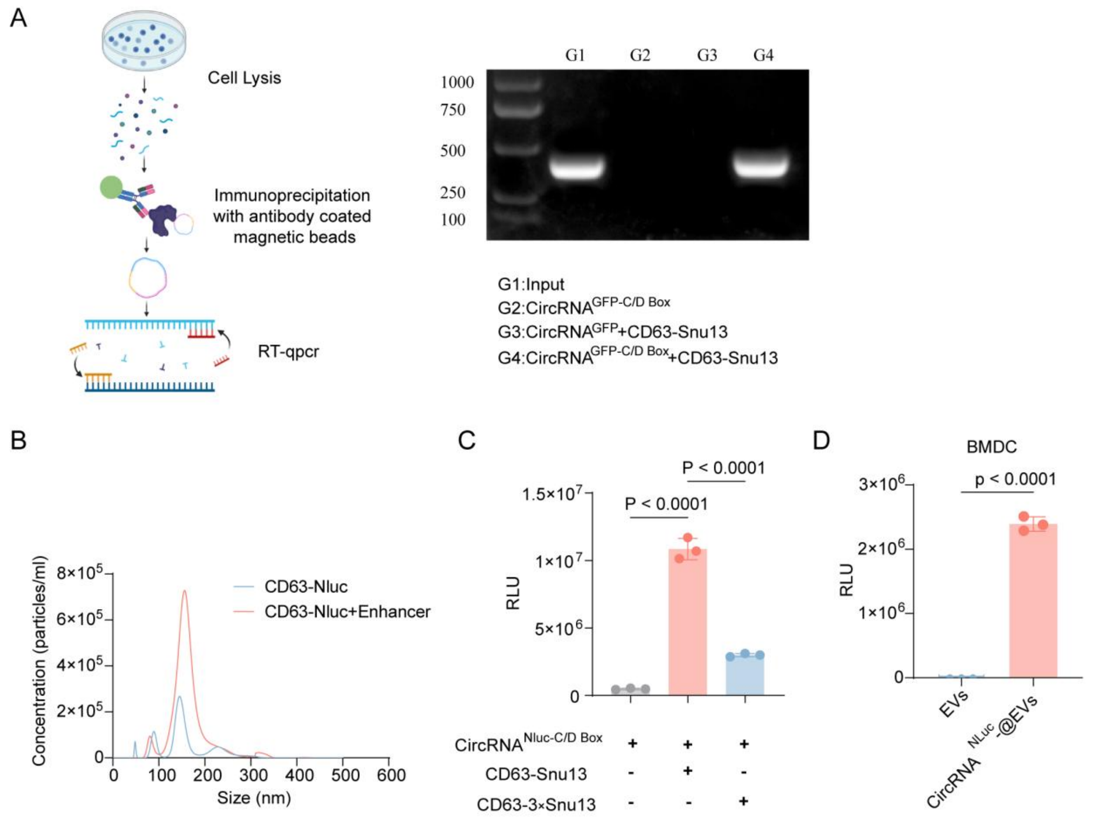


**fig. S2 Construction of an Engineered EV Platform for CircRNA Delivery.** (A) Diagram of the RNA immunoprecipitation assay used to assess CD63-Snu13 binding to circRNA. (B) Quantification of EVs derived from enhancer-transfected HEK-293T cells was performed using nanoparticle tracking analysis (NTA). (C) Impact of Snu13 tandem Repeats on circRNA Enrichment Efficiency. (D) The transfection efficacy of EV-delivered circRNA was evaluated in bone marrow-derived dendritic cells. Data are presented as mean ± SD. Statistical analysis in C and D were using one-way ANOVA with a Tukey multiple comparisons test.

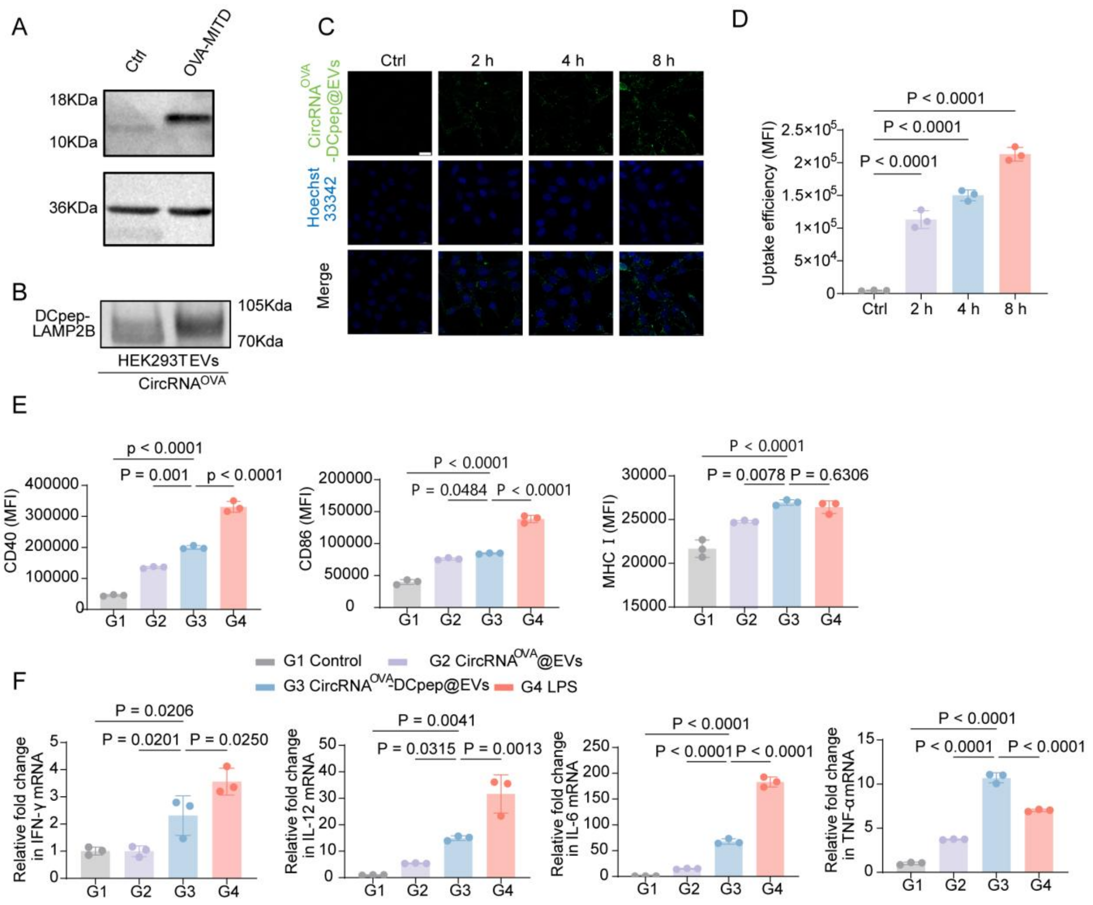


**fig. S3 Uptake and Maturation of Exosome-Delivered circRNA in BMDCs.** (A) OVA expression in HEK-293T cells following transfection with circRNA$^{OVA}$@EVs. (B) Surface display of the DCpep motif on engineered exosomes. (C-D) Cellular uptake of PKH67-labeled circRNAOVA-DCpep@EVs (green) by BMDCs over time. Nuclei were stained with DAPI (blue). Scale bar: 20 μm. Cellular uptake of PKH67-labeled circRNA$^{OVA}$-DCpep@EVs (green) by BMDCs over time. (D) MHC I-OVA complex expression on BMDCs treated with circRNA$^{OVA}$@EVs or circRNA$^{OVA}$-DCpep@EVs, quantified by flow cytometry. (E) Surface expression of maturation markers (CD40, CD86, MHC II) on CD11c$^{+}$ BMDCs after exposure to different formulations. (F) Evaluation of the innate immune activation of BMDC cells by the incubation with different formulations. Data are presented as mean ± SD. Statistical analysis were using one-way ANOVA with a Tukey multiple comparisons test.

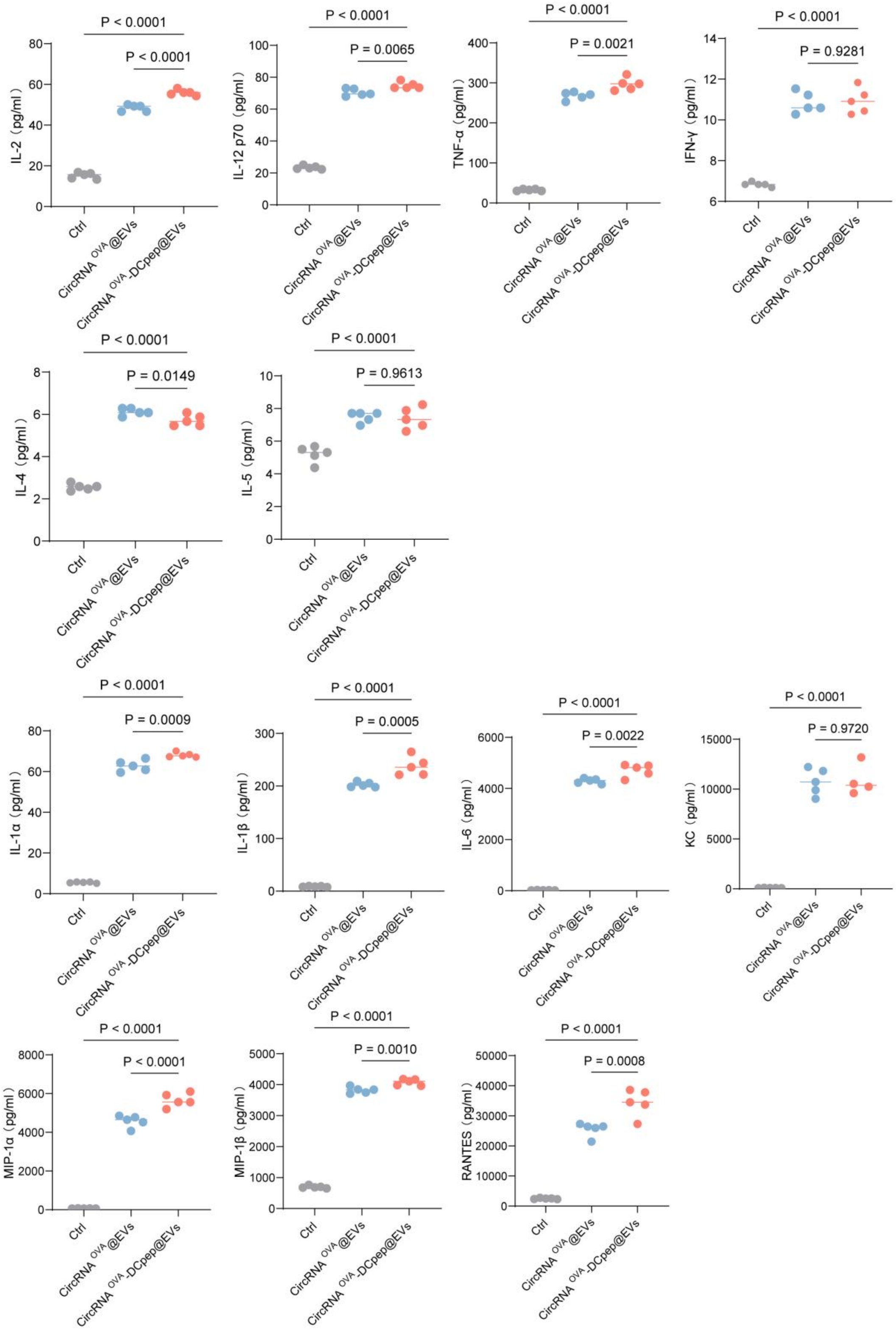


**fig. S4 The levels of IL-2, IL-12p70, TNF-α, IFN-γ, IL-4, IL-5, IL-1α, IL-1β, IL-6, KC, MIP-1α, MIP-1β and RANTES secreted by BMDCs after incubation with different formulations for 24 h (n = 5).** Data are presented as mean ± SD. Statistical analysis were using one-way ANOVA with a Tukey multiple comparisons test.

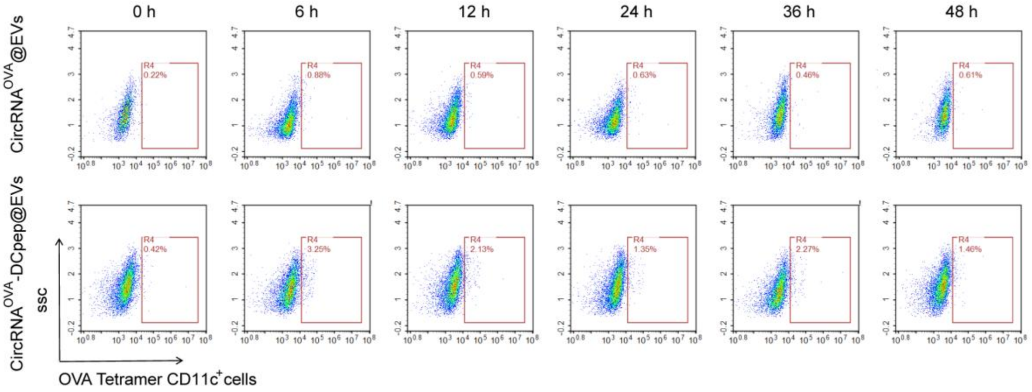


**fig. S5 Antigen presentation of Exosome-Delivered circRNA in BMDCs.**

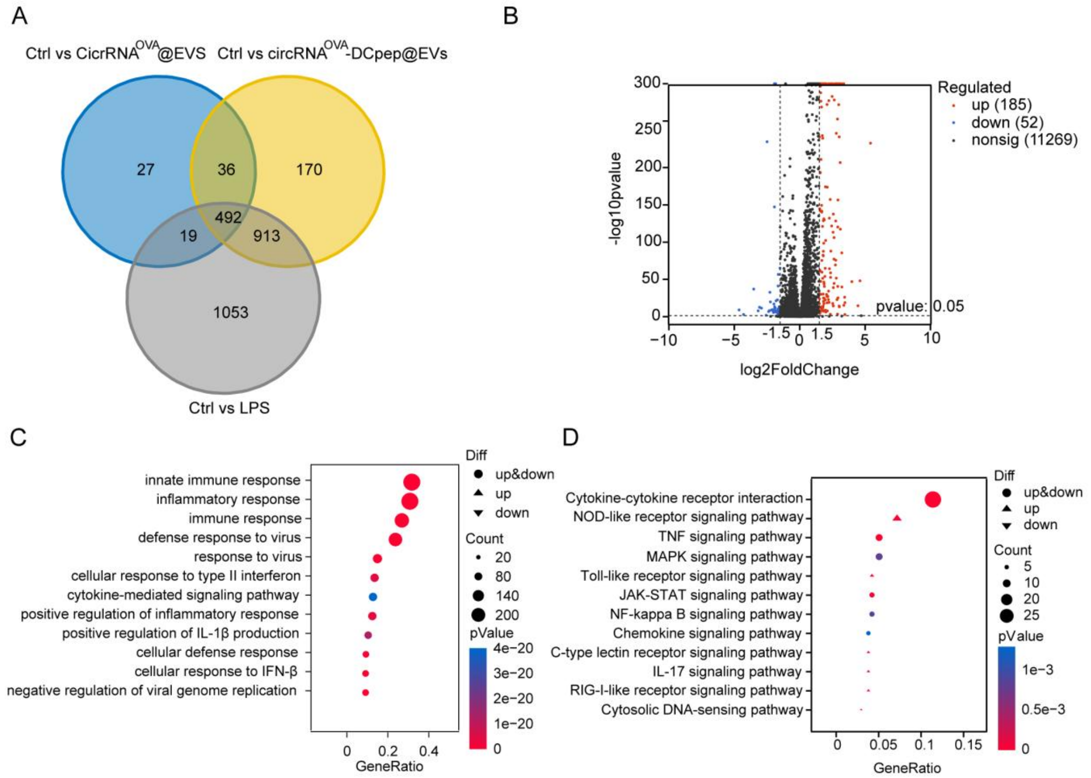


**fig. S6 RNA Sequencing reveals the mechanism of EVs-delivered circRNA Induced BMDC Maturation.** (A) Venn diagram illustrating overlapping and unique differentially expressed genes (DEGs) in BMDCs across treatment groups. (B) Volcano plot of DEGs between circRNA$^{OVA}$-DCpep@EVs and circRNA$^{OVA}$@EVs groups. (C) Gene Ontology (GO) enrichment analysis of up and down regulated pathways in circRNA$^{OVA}$-DCpep@EVs-treated BMDCs compared to the circRNA$^{OVA}$@EVs group. (D) KEGG pathway enrichment analysis of up and down regulated genes in the circRNA$^{OVA}$-DCpep@EVs group compared to the circRNA$^{OVA}$@EVs group.

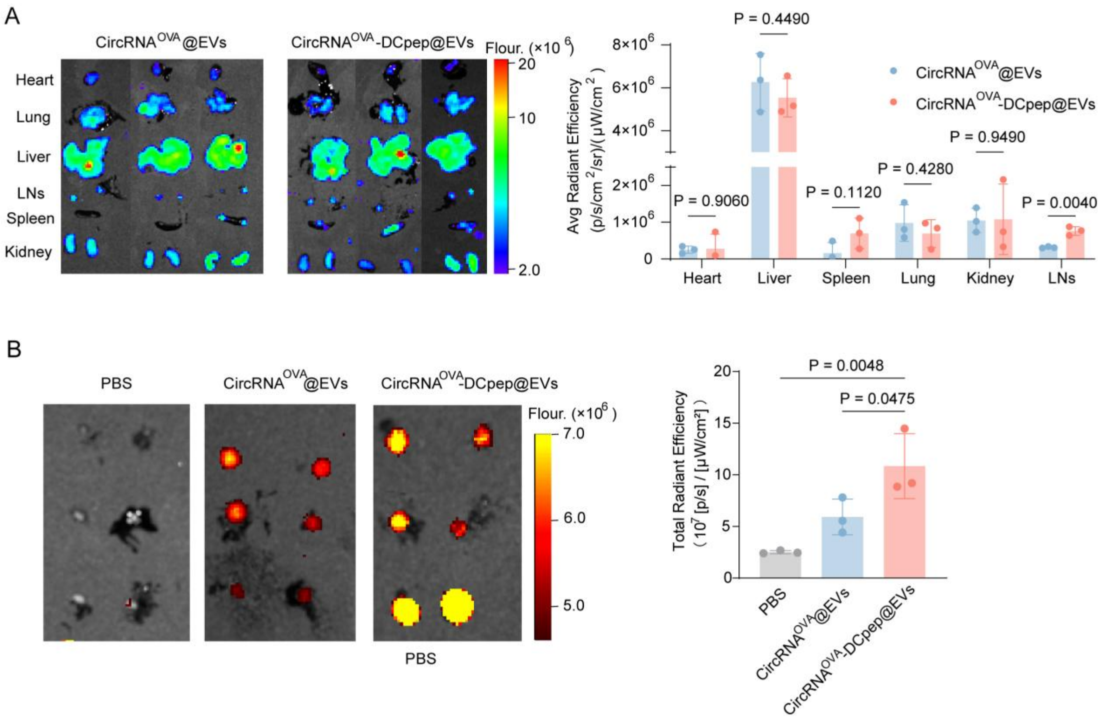


**fig. S7 IVIS image collected from mice treated with PBS, CircRNA$^{OVA}$@EVs or CircRNA$^{OVA}$-DCpep@EVs.** (A) Quantification of fluorescence intensity in major organs (heart, lung, liver, LNs, spleen, and kidney) collected from mice injected with PKH67-labelled EVs 24h post-vaccination. (B) IVIS image of LNs collected from mice treated with PBS, circRNA$^{OVA}$@EVs and circRNA$^{OVA}$-DCpep@EVs. Data are presented as mean ± SD. Statistical analysis were using one-way ANOVA with a Tukey multiple comparisons test.

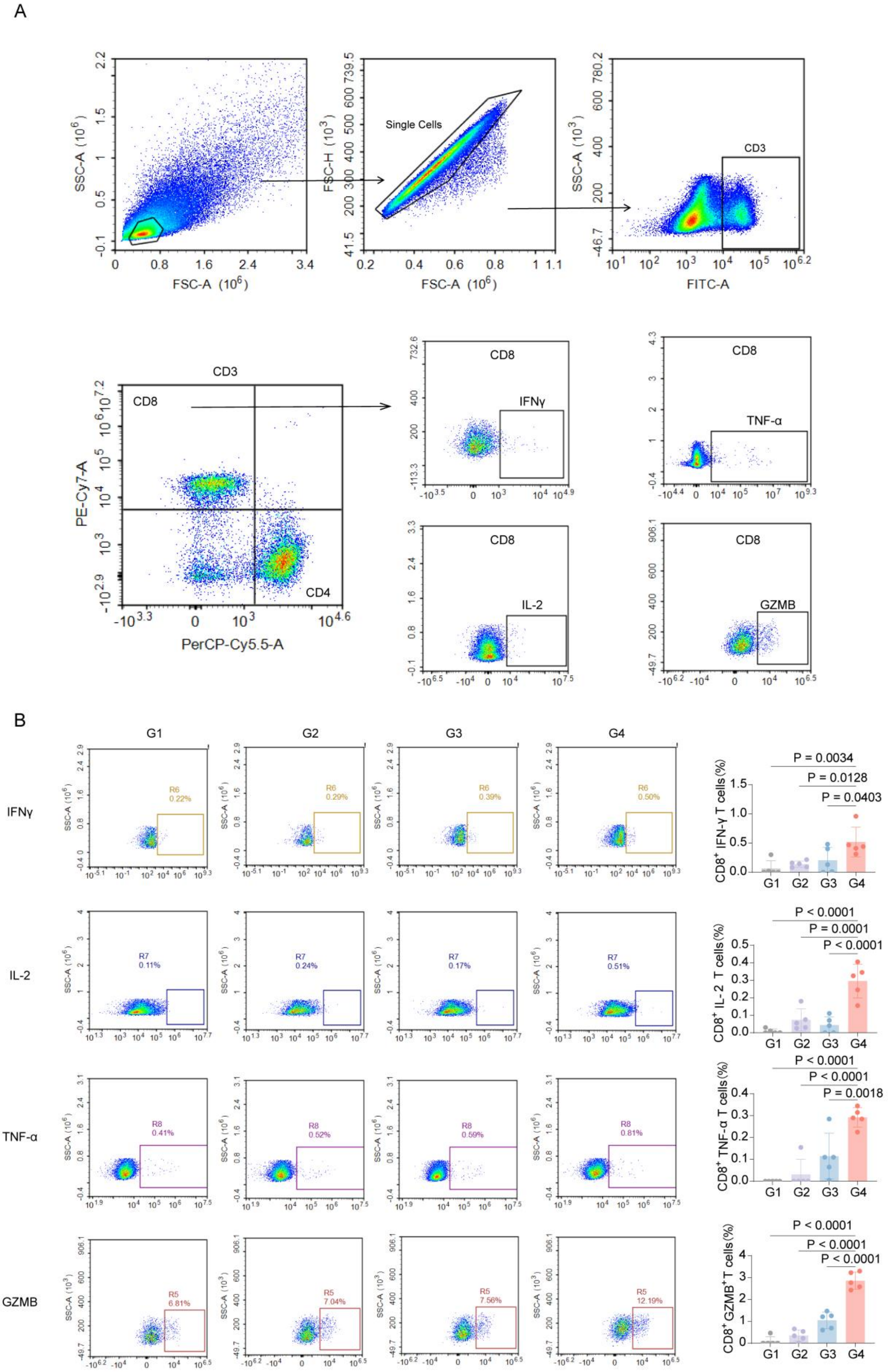


**fig. S8 Flow cytometric evaluation of splenic T cell populations shows the proportion of CD8$^{+}$ IFN-γ$^{+}$ T cells, CD8$^{+}$ IL-2$^{+}$ T cells, CD8$^{+}$ TNF-α$^{+}$ T cells and CD8$^{+}$ GZMB$^{+}$ T cells.** (A) Gating strategy for flow cytometric scatter plots to analyze the frequency of the splenic lymphocytes to secrete cytokines. (B)

Representative scatter plots of $CD8^+$ IFN-$\gamma^+$ T cells, $CD8^+$ IL-$2^+$ T cells, $CD8^+$ TNF-$\alpha^+$ T cells and $CD8^+$ $GZMB^+$ T cells in different groups.

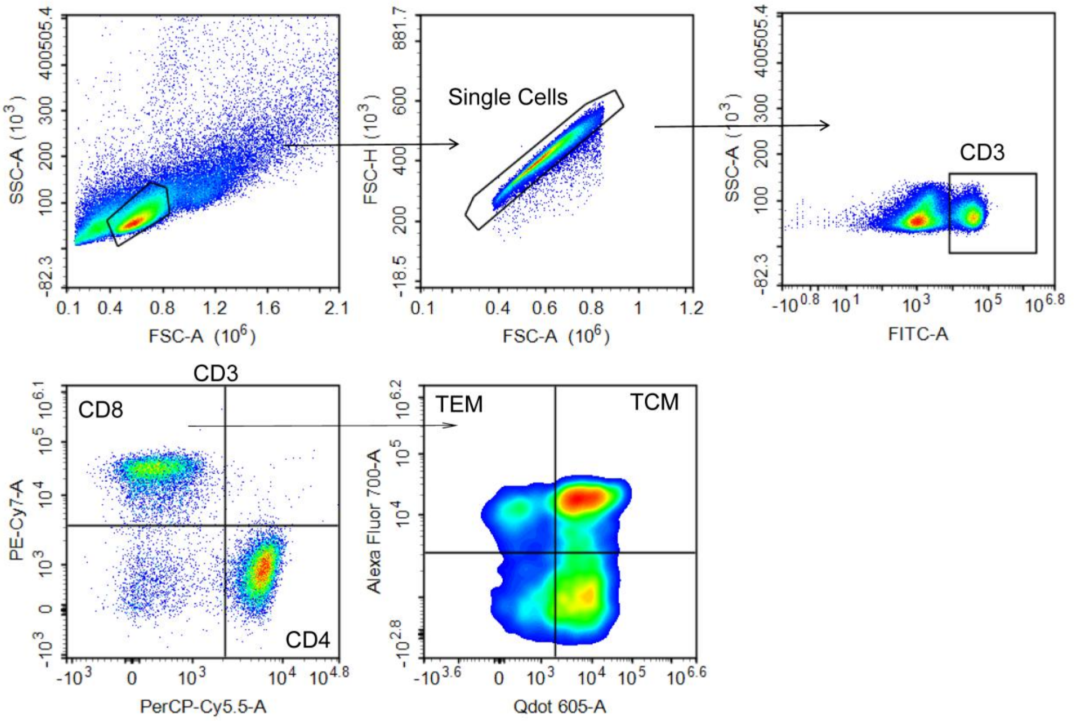


**fig. S9 Gating strategy for effector memory T cells (TEM, $CD3^{+}CD8^{+}CD44^{+}CD62L^{-}$) and central memory T cells (TCM, $CD3^{+}CD8^{+}CD44^{+}CD62L^{+}$) .**

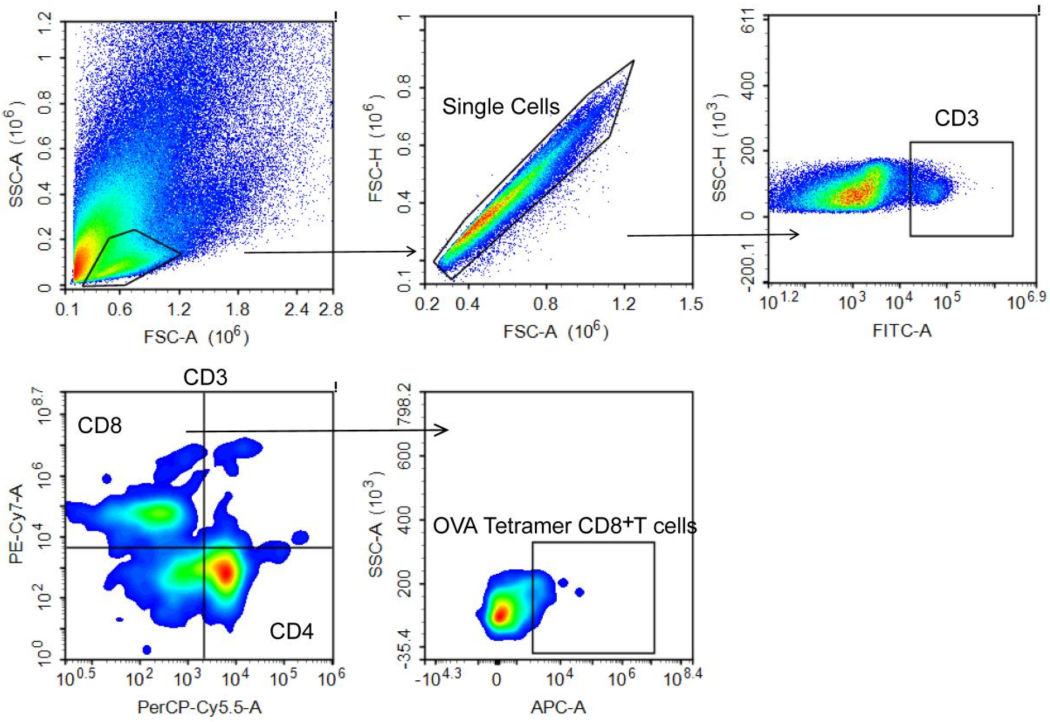


**fig. S10 Gating strategy for OVA Tetramer CD$^{+}$8 T cells.**

**Table S1. IRES Constructs Screened.**

| | |
|---|---|
| CircRNA expression vector<br>Structure Code: CMV Promoter, Spacer, Twister Ribozyme 5', Ligation Area 5', 5' UTR, IRES, GFP, C/D Box, 3' UTR, Ligation Area 3', Twister Ribozyme 3' | GTGATGCGGTTTTGGCAGTACATCAATGGGCGTGGATAGCGGTTTGACTCACGGGGATTTCC<br>AAGTCTCCACCCCATTGACGTCAATGGGAGTTTGTTTTGGCACCAAAATCAACGGGACTTTC<br>CAAAATGTCGTAACAACTCCGCCCCATTGACGCAAATGGGCGGTAGGCGTGTACGGTGGGA<br>GGTTTATATAAGCAGAGCT...(sapcer)...GGCCGCACTCGCCGGTCCCAAGCCCGGATAAAATGG<br>GAGGGGGCGGGAAACCGCCTAACCATGCCGAGTGCGGCCGCATATATGGATTCACAAAAAA<br>AACAAAAAAAAAAAAACAAAAAAAAAAAAACTAATTGACTAA...(IRES)...GCCACCATGGATA<br>TCGGCTTGGTGAGCAAGGGCGAGGAGCTGTTCACCGGGGTGGTGCCCATCCTGGTCGAGCT<br>GGACGGCGACGTAAACGGCCACAAGTTCAGCGTGTCCGGCGAGGGCGAGGGCGATGCCAC<br>CTACGGCAAGCTGACCCTGAAGTTCATCTGCACCACCGGCAAGCTGCCCGTGCCCTGGCCC<br>ACCCTCGTGACCACCCTGACCTACGGCGTGCAGTGCTTCAGCCGCTACCCCGACCACATGAA<br>GCAGCACGACTTCTTCAAGTCCGCCATGCCCGAAGGCTACGTCCAGGAGCGCACCATCTTCT<br>TCAAGGACGACGGCAACTACAAGACCCGCGCCGAGGTGAAGTTCGAGGGCGACACCCTGG<br>TGAACCGCATCGAGCTGAAGGGCATCGACTTCAAGGAGGACGGCAACATCCTGGGGCACAA<br>GCTGGAGTACAACTACAACAGCCACAACGTCTATATCATGGCCGACAAGCAGAAGAACGGC<br>ATCAAGGTGAACTTCAAGATCCGCCACAACATCGAGGACGGCAGCGTGCAGCTCGCCGACC<br>ACTACCAGCAGAACACCCCCATCGGCGACGGCCCCGTGCTGCTGCCCGACAACCACTACCT<br>GAGCACCCAGTCCGCCCTGAGCAAAGACCCCAACGAGAAGCGCGATCACATGGTCCTGCTG<br>GAGTTCGTGACCGCCGCCGGGATCACTCTCGGCATGGACGAGCTGTACAAGTAAGGGCGTG<br>ATGCGAAAGCTGACCCGGGCGTGATGCGAAAGCTGACCCGCTCTGACCGAAAGGCGTGATG<br>AGCGCTCTGACCGAAAGGCGTGATGAGCACCGGTGCTGGAGCCTCGGTGGCCATGCTTCTT<br>GCCCCTTGGGCCTCCCCCCAGCCCCTCCTCCCCTTCCTGCACCCGTACCCCCGTGGTCTTTGA<br>ATAAAGTCTGACCATGTCTATGTGGCCGCGGTCGGCGTGGACTGTAGAACACTGCCAATGCC<br>GGTCCCAAGCCCGGATAAAAGTGGAGGGTACAGTCCACGC |
| IRES_Hepatitis C Virus (HCV) | GCCAGCCCCCGATTGGGGGCGACACTCCACCATAGATCACTCCCCTGTGAGGAACTACTGTC<br>TTCACGCAGAAAGCGTCTAGCCATGGCGTTAGTATGAGTGTCGTGCAGCCTCCAGGACCCCC<br>CCTCCCGGGAGAGCCATAGTGGTCTGCGGAACCGGTGAGTACACCGGAATTGCCAGGACGA<br>CCGGGTCCTTTCTTGGATCAACCCGCTCAATGCCTGGAGATTTGGGCGTGCCCCCGCGAGAC<br>TGCTAGCCGAGTAGTGTTGGGTCGCGAAAGGCCTTGTGGTACTGCCTGATAGGGTGCTTGCG<br>AGTGCCCCGGGAGGTCTCGTAGACCGTGCACCATGAGCACGAATCCTAAACCTCAAAGAAA<br>AACC |
| IRES_Enterovirus 71 (EV71) | TTAAAACAGCTGTGGGTTGTCACCCACCCACAGGGTCCACTGGGCGCTAGTACACTGGTATC<br>TCGGTACCTTTGTACGCCTGTTTTATACCCCCTCCCTGATTTGCAACTTAGAAGCAACGCAAA<br>CCAGATCAATAGTAGGTGTGACATACCAGTCGCATCTTGATCAAGCACTTCTGTATCCCCGGA<br>CCGAGTATCAATAGACTGTGCACACGGTTGAAGGAGAAAACGTCCGTTACCCGGCTAACTAC<br>TTCGAGAAGCCTAGTAACGCCATTGAAGTTGCAGAGTGTTTCGCTCAGCACTCCCCCCGTGT<br>AGATCAGGTCGATGAGTCACCGCATTCCCCACGGGCGACCGTGGCGGTGGCTGCGTTGGCG<br>GCCTGCCTATGGGGTAACCCATAGGACGCTCTAATACGGACATGGCGTGAAGAGTCTATTGA<br>GCTAGTTAGTAGTCCTCCGGCCCCTGAATGCGGCTAATCCTAACTGCGGAGCACATACCCTTA<br>ATCCAAAGGGCAGTGTGTCGTAACGGGCAACTCTGCAGCGGAACCGACTACTTTGGGTGTC<br>CGTGTTTCTTTTTATTCTTGTATTGGCTGCTTATGGTGACAATTAAAGAATTGTTACCATATAGC<br>TATTGGATTGGCCATCCAGTGTCAAACAGAGCTATTGTATATCTCTTTGTTGGATTCACACCTC<br>TCACTCTTGAAACGTTACACACCCTCAATTACATTATACTGCTGAACACGAAGCG |

| | |
|---|---|
| IRES_Encephalomyocarditis Virus_6A (ECMV) | CCCCTCTCCCTCCCCCCCCCCTAACGTTACTGGCCGAAGCCGCTTGGAATAAGGCCGGTGTG<br>CGTTTGTCTATATGTTATTTTCCACCATATTGCCGTCTTTTGGCAATGTGAGGGCCCGGAAACC<br>TGGCCCTGTCTTCTTGACGAGCATTCCTAGGGGTCTTTCCCCTCTCGCCAAAGGAATGCAAG<br>GTCTGTTGAATGTCGTGAAGGAAGCAGTTCCTCTGGAAGCTTCTTGAAGACAAACAACGTC<br>TGTAGCGACCCTTTGCAGGCAGCGGAACCCCCCACCTGGCGACAGGTGCCTCTGCGGCCAA<br>AAGCCACGTGTATAAGATACACCTGCAAAGGCGGCACAACCCCAGTGCCACGTTGTGAGTT<br>GGATAGTTGTGGAAAGAGTCAAATGGCTCTCCTCAAGCGTATTCAACAAGGGGCTGAAGGAT<br>GCCCAGAAGGTACCCCATTGTATGGGATCTGATCTGGGGCCTCGGTGCACATGCTTTACATGT<br>GTTTAGTCGAGGTTAAAAAACGTCTAGGCCCCCCGAACCACGGGGACGTGGTTTTCCTTTGA<br>AAAACACGATGATAATATGGCCACAACC |
| IRES_Coxsackievirus B3 (CVB3) | TTAAAAGAATTCCAGCCTGTGGGTTGATCCCACCCACAGGCCCATTGGGCGCTAGCACTCTG<br>GTATCACGGTACCTTTGTGCGCCTGTTTTATACCCCCTCCCCCAACTGTAACTTAGAAGTAAC<br>ACACACCGATCAACAGTCAGCGTGGCACACCAGCCACGTTTTGATCAAGCACTTCTGTTACC<br>CCGGACTGAGTATCAATAGACTGCTCACGCGGTTGAAGGAGAAAGCGTTCGTTATCCGGCCA<br>ACTACTTCGAAAAACCTAGTAACACCGTGGAAGTTGCAGAGTGTTTCGCTCAGCACTACCCC<br>AGTGTAGATCAGGTCGATGAGTCACCGCATTCCCCACGGGCGACCGTGGCGGTGGCTGCGTT<br>GGCGGCCTGCCCATGGGGAAACCCATGGGACGCTCTAATACAGACATGGTGCGAAGAGTCTA<br>TTGAGCTAGTTGGTAGTCCTCCGGCCCCTGAATGCGGCTAATCCTAACTGCGGAGCACACAC<br>CCTCAAGCCAGAGGGCAGTGTGTCGTAACGGGCAACTCTGCAGCGGAACCGACTACTTTGG<br>GTGTCCGTGTTTCATTTTATTCCTATACTGGCTGCTTATGGTGACAATTGAGAGATCGTTACCA<br>TATAGCTATTGGATTGGCCATCCGGTGACTAATAGAGCTATTATATATCCCTTTGTTGGGTTTAT<br>ACCACTTAGCTTGAAAGAGGTTAAAACATTACAATTCATTGTTAAGTTGAATACAGCAAA |
| IRES_Human rhinovirus B3 (HRVB3) | TTAAAACAGCGGATGGGTACCCCACCATCCGACCCACTGGGTGTAGTACTCTGGTACTTCGT<br>ACCTTTGTACGCCTGTTCTTCCCATTGTACCCTTCCTGAACTTCCAACCCAAGTAACGTTAGA<br>AGCTCAACATTTAGTACAACAGGAAGCACCACATCCAGTGGTGTTTAGTACAAGCACTTCTG<br>TTTCCCCGGAGCGAGGTATAGGCTGTACCCACTGCCAAAAACCTTTAACCGTTATCCGCCAA<br>CCAACTACGTAAAAGCTAGTAGTATTATGTTTTTAACTAGGCGTTCGATCAGGTGGATTTCCC<br>CTCCACTAGTTTGGTCGATGAGGCTAGGAATTCCCCACGGGTGACCGTGTCCTAGCCTGCGT<br>GGCGGCCAACCCAGCCCACTCACTATTTGTTTTCGCGCCCAGTTGCAAAAAGTGTCGGGGCT<br>GGGACGCCTTTTTATAGACATGGTGTGAAGACTCGCATGTGCTTGGTTGTGATTCCTCCGGCC<br>CCTGAATGCGGCTAACCTTAACCCTGGAGCCTTGTGTCACAAACCAGTGATGATAAGGTCGT<br>AATGAGCAATTCCGGGACGGGACCGACTACTTTGGGTGTCCGTGTTTCTTATTTTTCTTATTAT<br>TGTCTTATGGTCACAGCATATATATAACATATACTGTGATC |

**Table S2. Translation originals Screened.**

| | |
|---|---|
| #2 | GTGATGCGGTTTTGGCAGTACATCAATGGGCGTGGATAGCGGTTTGACTCACGGGGATTTCC |

| | |
|---|---|
| Structure Code: CMV Promoter, Spacer, Twister Ribozyme 5’, Ligation Area 5’, 5’ UTR, IRES, GFP, C/D Box, 3’ UTR, Ligation Area 3’, Twister Ribozyme 3’, WPRE | AAGTCTCCACCCCATTGACGTCAATGGGAGTTTGTTTTGGCACCAAAATCAACGGGACTTTC<br>CAAAATGTCGTAACAACTCCGCCCCATTGACGCAAATGGGCGGTAGGCGTGTACGGTGGGA<br>GGTTTATATAAGCAGAGCT...(sapcer)...GGCCGCACTCGCCGGTCCCAAGCCCGGATAAAATGG<br>GAGGGGGCGGGAAACCGCCTAACCATGCCGAGTGCGGCCGCATATATGGATTCACAAAAAA<br>AACAAAAAAAAAAAAACAAAAAAAAAAAAACTAATTGACTAA...(IRES)...GCCACCATGGATA<br>TCGGCTTGGTGAGCAAGGGCGAGGAGCTGTTCACCGGGGTGGTGCCCATCCTGGTCGAGCT<br>GGACGGCGACGTAAACGGCCACAAGTTCAGCGTGTCCGGCGAGGGCGAGGGCGATGCCAC<br>CTACGGCAAGCTGACCCTGAAGTTCATCTGCACCACCGGCAAGCTGCCCGTGCCCTGGCCC<br>ACCCTCGTGACCACCCTGACCTACGGCGTGCAGTGCTTCAGCCGCTACCCCGACCACATGAA<br>GCAGCACGACTTCTTCAAGTCCGCCATGCCCGAAGGCTACGTCCAGGAGCGCACCATCTTCT<br>TCAAGGACGACGGCAACTACAAGACCCGCGCCGAGGTGAAGTTCGAGGGCGACACCCTGG<br>TGAACCGCATCGAGCTGAAGGGCATCGACTTCAAGGAGGACGGCAACATCCTGGGGCACAA<br>GCTGGAGTACAACTACAACAGCCACAACGTCTATATCATGGCCGACAAGCAGAAGAACGGC<br>ATCAAGGTGAACTTCAAGATCCGCCACAACATCGAGGACGGCAGCGTGCAGCTCGCCGACC<br>ACTACCAGCAGAACACCCCCATCGGCGACGGCCCCGTGCTGCTGCCCGACAACCACTACCT<br>GAGCACCCAGTCCGCCCTGAGCAAAGACCCCAACGAGAAGCGCGATCACATGGTCCTGCTG<br>GAGTTCGTGACCGCCGCCGGGATCACTCTCGGCATGGACGAGCTGTACAAGTAAGGGCGTG<br>ATGCGAAAGCTGACCCGGGCGTGATGCGAAAGCTGACCCGCTCTGACCGAAAGGCGTGATG<br>AGCGCTCTGACCGAAAGGCGTGATGAGCACCGGTGCTGGAGCCTCGGTGGCCATGCTTCTT<br>GCCCCTTGGGCCTCCCCCCAGCCCCTCCTCCCCTTCCTGCACCCGTACCCCCGTGGTCTTTGA<br>ATAAAGTCTGACCATGTCTATGTGGCCGCGGTCGGCGTGGACTGTAGAACACTGCCAATGCC<br>GGTCCCAAGCCCGGATAAAAGTGGAGGGTACAGTCCACGCAATCAACCTCTGGATTACAAA<br>ATTTGTGAAAGATTGACTGGTATTCTTAACTATGTTGCTCCTTTTACGCTATGTGGATACGCTG<br>CTTTAATGCCTTTGTATCATGCTATTGCTTCCCGTATGGCTTTCATTTTCTCCTCCTTGTATAAAT<br>CCTGGTTGCTGTCTCTTTATGAGGAGTTGTGGCCCGTTGTCAGGCAACGTGGCGTGGTGTGC<br>ACTGTGTTTGCTGACGCAACCCCCACTGGTTGGGGCATTGCCACCACCTGTCAGCTCCTTTC<br>CGGGACTTTCGCTTTCCCCCTCCCTATTGCCACGGCGGAACTCATCGCCGCCTGCCTTGCCCG<br>CTGCTGGACAGGGGCTCGGCTGTTGGGCACTGACAATTCCGTGGTGTTGTCGGGGAAATCAT<br>CGTCCTTTCCTTGGCTGCTCGCCTGTGTTGCCACCTGGATTCTGCGCGGGACGTCCTTCTGCT<br>ACGTCCCTTCGGCCCTCAATCCAGCGGACCTTCCTTCCCGCGGCCTGCTGCCGGCTCTGCGG<br>CCTCTTCCGCGTCTTCGCCTTCGCCCTCAGACGAGTCGGATCTCCCTTTGGGCCGCCTCCCCG<br>C |
| #3<br>Structure Code: 5’LTR, CMV Promoter, Spacer, Twister Ribozyme 5’, Ligation Area 5’, 5’ UTR, IRES, GFP, C/D Box, 3’ UTR, Ligation Area 3’, Twister Ribozyme 3’, WPRE, 3’LTR | GGGTCTCTCTGGTTAGACCAGATCTGAGCCTGGGAGCTCTCTGGCTAACTAGGGAACCCACT<br>GCTTAAGCCTCAATAAAGCTTGCCTTGAGTGCTTCAAGTAGTGTGTGCCCGTCTGTTGTGTG<br>ACTCTGGTAACTAGAGATCCCTCAGACCCTTTTAGTCAGTGTGGAAAATCTCTAGCAGTGAT<br>GCGGTTTTGGCAGTACATCAATGGGCGTGGATAGCGGTTTGACTCACGGGGATTTCCAAGTC<br>TCCACCCCATTGACGTCAATGGGAGTTTGTTTTGGCACCAAAATCAACGGGACTTTCCAAAA<br>TGTCGTAACAACTCCGCCCCATTGACGCAAATGGGCGGTAGGCGTGTACGGTGGGAGGTTTA<br>TATAAGCAGAGCT...(sapcer)...GGCCGCACTCGCCGGTCCCAAGCCCGGATAAAATGGGAGGG<br>GGCGGGAAACCGCCTAACCATGCCGAGTGCGGCCGCATATATGGATTCACAAAAAAAACAA<br>AAAAAAAAACAAAAAAAAAAAAACTAATTGACTAA...(IRES)...GCCACCATGGATATCGGCT<br>TGGTGAGCAAGGGCGAGGAGCTGTTCACCGGGGTGGTGCCCATCCTGGTCGAGCTGGACGG<br>CGACGTAAACGGCCACAAGTTCAGCGTGTCCGGCGAGGGCGAGGGCGATGCCACCTACGGC<br>AAGCTGACCCTGAAGTTCATCTGCACCACCGGCAAGCTGCCCGTGCCCTGGCCCACCCTCGT |

| | |
|---|---|
| | GACCACCCTGACCTACGGCGTGCAGTGCTTCAGCCGCTACCCCGACCACATGAAGCAGCAC<br>GACTTCTTCAAGTCCGCCATGCCCGAAGGCTACGTCCAGGAGCGCACCATCTTCTTCAAGGA<br>CGACGGCAACTACAAGACCCGCGCCGAGGTGAAGTTCGAGGGCGACACCCTGGTGAACCG<br>CATCGAGCTGAAGGGCATCGACTTCAAGGAGGACGGCAACATCCTGGGGCACAAGCTGGAG<br>TACAACTACAACAGCCACAACGTCTATATCATGGCCGACAAGCAGAAGAACGGCATCAAGG<br>TGAACTTCAAGATCCGCCACAACATCGAGGACGGCAGCGTGCAGCTCGCCGACCACTACCA<br>GCAGAACACCCCCATCGGCGACGGCCCCGTGCTGCTGCCCGACAACCACTACCTGAGCACC<br>CAGTCCGCCCTGAGCAAAGACCCCAACGAGAAGCGCGATCACATGGTCCTGCTGGAGTTCG<br>TGACCGCCGCCGGGATCACTCTCGGCATGGACGAGCTGTACAAGTAAGGGCGTGATGCGAA<br>AGCTGACCCGGGCGTGATGCGAAAGCTGACCCGCTCTGACCGAAAGGCGTGATGAGCGCTC<br>TGACCGAAAGGCGTGATGAGCACCGGTGCTGGAGCCTCGGTGGCCATGCTTCTTGCCCCTTG<br>GGCCTCCCCCCAGCCCCTCCTCCCCTTCCTGCACCCGTACCCCCGTGGTCTTTGAATAAAGTC<br>TGACCATGTCTATGTGGCCGCGGTCGGCGTGGACTGTAGAACACTGCCAATGCCGGTCCCAA<br>GCCCGGATAAAAGTGGAGGGTACAGTCCACGCAATCAACCTCTGGATTACAAAATTTGTGAA<br>AGATTGACTGGTATTCTTAACTATGTTGCTCCTTTTACGCTATGTGGATACGCTGCTTTAATGC<br>CTTTGTATCATGCTATTGCTTCCCGTATGGCTTTCATTTTCTCCTCCTTGTATAAATCCTGGTTG<br>CTGTCTCTTTATGAGGAGTTGTGGCCCGTTGTCAGGCAACGTGGCGTGGTGTGCACTGTGTT<br>TGCTGACGCAACCCCCACTGGTTGGGGCATTGCCACCACCTGTCAGCTCCTTTCCGGGACTT<br>TCGCTTTCCCCCTCCCTATTGCCACGGCGGAACTCATCGCCGCCTGCCTTGCCCGCTGCTGGA<br>CAGGGGCTCGGCTGTTGGGCACTGACAATTCCGTGGTGTTGTCGGGGAAATCATCGTCCTTT<br>CCTTGGCTGCTCGCCTGTGTTGCCACCTGGATTCTGCGCGGGACGTCCTTCTGCTACGTCCCT<br>TCGGCCCTCAATCCAGCGGACCTTCCTTCCCGCGGCCTGCTGCCGGCTCTGCGGCCTCTTCC<br>GCGTCTTCGCCTTCGCCCTCAGACGAGTCGGATCTCCCTTTGGGCCGCCTCCCCGCTGGAAG<br>GGCTAATTCACTCCCAACGAAAATAAGATCTGCTTTTTGCTTGTACTGGGTCTCTCTGGTTAG<br>ACCAGATCTGAGCCTGGGAGCTCTCTGGCTAACTAGGGAACCCACTGCTTAAGCCTCAATAA<br>AGCTTGCCTTGAGTGCTTCAAGTAGTGTGTGCCCGTCTGTTGTGTGACTCTGGTAACTAGAG<br>ATCCCTCAGACCCTTTTAGTCAGTGTGGAAAATCTCTAGCA |

**Table S3. The primer sequence for qPCR.**

| Primer name | The primer sequence (5'-3') |
|---|---|

| | |
|---|---|
| GFP-qPCR-F (Convergent primer) | CGTGACCACCCTGACCT |
| GFP-qPCR-R (Convergent primer) | CACCTTGATGCCGTTCTT |
| TornadoOpti-CJ-F (Divergent primer) | CCCCGTGGTCTTTGA |
| TornadoOpti-CJ-R (Divergent primer) | GTTGGTTGGCGGATAA |
| qm-IFN-γ-F | ACAGCAAGGCGAAAAAGGATG |
| qm-IFN-γ-R | TGGTGGACCACTCGGATGA |
| qm-IL-12β-F | TGGTTTGCCATCGTTTTGCTG |
| qm-IL-12β-R | ACAGGTGAGGTTCACTGTTTCT |
| qm-IL-6-F | TAGTCCTTCCTACCCCAATTTCC |
| qm-IL-6-R | TTGGTCCTTAGCCACTCCTTC |
| qm-TNF-a-F | CCCTCACACTCAGATCATCTTCT |
| qm-TNF-a-F | GCTACGACGTGGGCTACAG |
| qh-GAPDH-F | GGAGCGAGATCCCTCCAAAAT |
| qh-GAPDH-R | GGCTGTTGTCATACTTCTCATGG |
| Rev-qhGAPDH-F | CCCCTCCTCACAGTTGCCATG |
| Rev-qhGAPDH-R | CGCAGGGTTAGTCACCGGCA |
| qm-GAPDH-F | AGGTCGGTGTGAACGGATTTG |
| qm-GAPDH-R | TGTAGACCATGTAGTTGAGGTCA |
